# Brainwide representation of navigation

**DOI:** 10.64898/2026.08.29.747979

**Authors:** Enny H. van Beest, Bex Terry, George Booth, Kenneth D. Harris, Matteo Carandini

## Abstract

Tracking one’s position in the environment during navigation requires integrating multiple signals into a spatial representation. This process is thought to occur primarily in the hippocampal formation, but evidence for it has also been observed in other regions. To investigate it across the brain, we recorded from over 20,000 neurons in mice navigating a virtual corridor that decoupled position from correlated signals. Spatial position was encoded in every region, and was more common in neurons tuned to visual landmarks. Most neurons brainwide were also modulated by running speed, likely reflecting arousal, and many by rewards. The hippocampal formation represented spatial position slightly more uniformly than other regions, but less precisely than visual cortex. Spatial position and other navigational signals are thus represented widely across the brain.

## Introduction

Since the discovery of place cells in the hippocampus (*1*) signals related to the body’s position in the environment have been claimed in a variety of cortical regions. Spatial representations have been found not only in other regions of the hippocampal formation (*2–9*) but also in parietal (*10–15*) and retrosplenial cortex (*10*, *14*, *16–18*). Since then, they have been claimed in many other cortical regions, such as medial prefrontal (*19*) and orbitofrontal (*20*, *21*) areas and even primary sensory and motor areas (*14*, *22–29*).

However, these potential spatial representations have been typically observed in isolation, using uneven approaches to distinguish spatial position from other correlated signals. Even in the hippocampus, where spatial representations are most established, neurons are modulated by many correlated signals, including heading direction (*1*, *30–33*), running speed (*33–35*), elapsed time (*36*, *37*), reward expectation (*38–40*), visual cues (*41–47*), and, in some conditions, auditory cues (*48–50*). These signals may be mixed with representations of space also in other regions, and the mixture may depend on the region. For example, some cortical areas are more visual than others (*51*, *52*), and different areas tend to have different correlates with running speed (*53*, *54*). Efforts to distinguish these correlated signals from spatial representations have been uneven. For instance, studies that controlled for the effects of running speed only investigated a few regions (*12*, *24*, *25*, *55*), whereas a study that focused on a wider set of regions could not control for the effects of running speed and sensory inputs (*14*).

It also remains unclear whether spatial representations extend beyond the cortex. Potential encoding of space has been claimed in regions such as the striatum (*56*, *57*), the claustrum (*58*), the lateral septum (*59–61*), and the cerebellum (*62*). Once again, however, these spatial representations were measured with different approaches; it is unclear whether they are all distinct from representations of other correlated signals.

To investigate spatial representations across the brain, we surveyed the activity of 22,965 neurons while mice navigated a virtual corridor that decoupled position from other correlated signals. We used Neuropixels probes to record from the midbrain, thalamus, striatum, neocortex, and hippocampal formation while head-fixed mice traversed a virtual corridor defined by audiovisual landmarks. Virtual reality allows the study of navigation during head fixation (*15*, *63*, *64*) while disentangling spatial position from other navigational signals such as running speed (*46*, *65*, *66*) and visual input (*24*, *25*, *66*). We could thus manipulate the corridor to decouple spatial position from the presence of auditory and visual landmarks and from variables related to motion and reward, allowing us to distinguish the impact of these signals on neural activity.

The results revealed a highly distributed organization of navigation signals across the brain. We found navigational signals related to position, motion, sensation, and reward not only across cortical regions but also widely across the rest of the forebrain and in the midbrain, often intermixed within the same neurons. Navigational signals were anchored to visual but not auditory landmarks and were more common in neurons which were tuned to sensory (and especially visual) stimuli when assessed during passive stimulus presentation not involving virtual reality. Activity in every brain region correlated strongly with running speed, potentially related to variations in arousal. The hippocampal formation did not have a higher fraction of neurons encoding spatial position than other regions. Its spatial representation was slightly more uniform than in other regions, but less precise than in the visual cortex.

## Results

We recorded activity with Neuropixels probes while mice navigated a virtual corridor with two identical segments spaced 40 cm apart (**Figure 1a,b**). Each segment contained a pair of audiovisual landmarks (called A and B, **Figure 1c**). To ensure that the audiovisual inputs at any position between 5 and 100 cm were identical to those 40 cm away, the corridor represented the segment multiple times (ABABABAB…). However, when the mouse progressed 100 cm into the corridor (in the middle of the last A in ABABA), the screen turned gray for 2-3 s, and the mouse was placed at the beginning of the corridor. To provide motivation, mice were water-limited and received a probabilistic water reward at a fixed location (typically 75 cm or 95 cm, depending on the mouse). Across trials we jointly varied the visual contrast and sound intensity of both landmarks, and we varied the gain between the running wheel and the virtual corridor (**Figure 1b**). We recorded 162,252 units across the forebrain and midbrain (**Figure 1d**), of which 22,965 were unique and well-isolated neurons (*67*, *68*) (**Figure S1**). Anatomical positions were assigned based on histology (**Figure S2**). For statistical analyses, we only included brain regions with at least 25 neurons in at least two mice, leaving 22,014 neurons across 38 regions in 28 mice (**Table S2**). For some analyses and visualizations, we aggregated brain regions in 8 groups: hippocampal formation, visual cortex, somatosensory cortex, motor cortex, other cortex (including retrosplenial and auditory cortex), basal ganglia (dorsal striatum and medial pallidum), thalamus, and midbrain (including medial reticular nucleus and motor-related superior colliculus).

**Figure 1.**
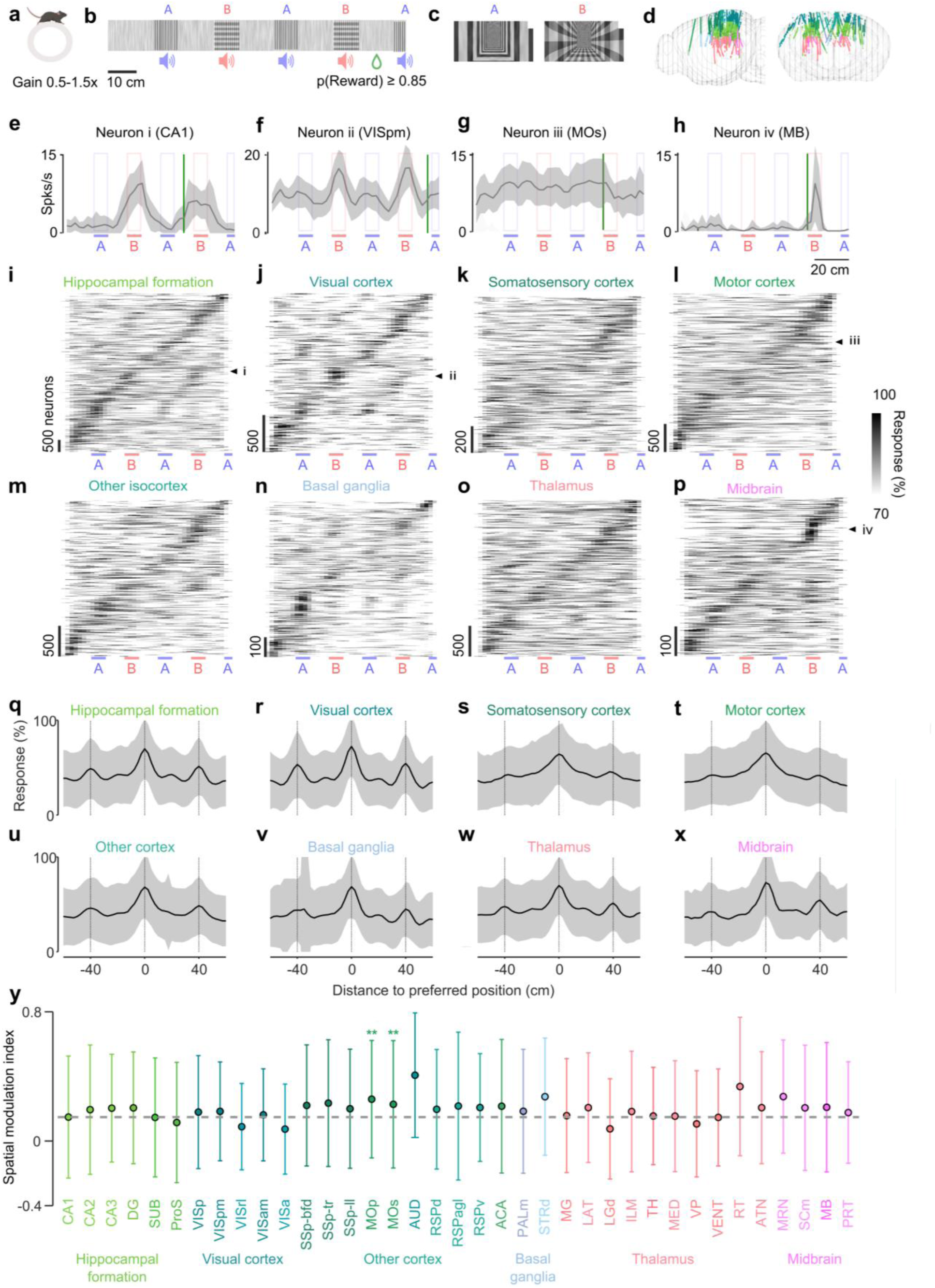
Brainwide navigational signals. **a)** The animal traverses the virtual corridor on a running wheel with random gain between 0.5 and 1.5x. **b)** The corridor contains audiovisual landmarks A and B, 20 cm apart, in the order ABABA. Visual landmarks are a grating (A) and a plaid (B) with variable contrast over a noise background. Auditory landmarks (when present) are pure tones at 11 kHz and 17 kHz, amplitude-modulated at 5.0 Hz and 7.1 Hz, ∼65 dB at the landmark position, and progressively weaker at more distant positions. A probabilistic reward (p≥0.85) is delivered towards the end of the corridor. **c)** Views of the two visual landmarks in the corridor, at full contrast. **d)** Coronal and sagittal view of Neuropixels trajectories, with colors adapted from the Allen Mouse Brain Common Coordinate Framework. **e)** Activity of an example neuron in hippocampal area CA1 (mean ± s.d. across full-contrast trials) as a function of position in the corridor. Reward position is indicated by the vertical green bar. **f-h)** Same, for example neurons in posteromedial visual cortex (VISpm), secondary motor cortex (MOs) and midbrain (MB). **i)** Activity of all neurons in the hippocampal formation, normalized, sorted by preferred position on held-out trials. Arrow points to the example neuron in e. Note that reward position was not perfectly consistent across neurons (the effect of reward is analyzed in Figure 6). **j-p)** Same, for 7 other groups of regions: visual cortex, somatosensory cortex, motor cortex, other cortex (retrosplenial, anterior cingulate, and auditory cortex), basal ganglia (striatum, pallidum, and lateral septum), thalamus, and midbrain. **q-x**) Normalized activity (mean ± s.d. % across neurons) as a function of distance to preferred position (measured in held-out trials) for the same brain groups, measured with visual landmarks at full contrast. 20.5% of neurons with unreliable responses were excluded from these analyses (see Methods). Lines indicate positions that are sensory identical to the preferred position. **y)** Spatial modulation index (i.e. difference between preferred and average of sensory identical positions divided by the sum, mean ± s.d. across included neurons). **, p<0.01 for post-hoc comparisons of brain region versus CA1, significant fixed effect of brain region in LME. Dashed line indicates mean value for CA1, for comparison.

### Brainwide navigational signals

Neurons in all recorded regions carried signals related to position, typically tiling the entire corridor. Many neurons showed spatially selective responses along the corridor, often with two peaks separated by 40 cm, the distance between the two identical corridor segments (e.g. **Figure 1e-f**). Other neurons had broader response fields with reduced activity after the reward position (**Figure 1g**), or with a peak just after the reward position (**Figure 1h**). Remarkably, neurons from all regions had spatial firing correlates that tiled the corridor (**Figure 1i,j**), including subcortical regions such as thalamus, basal ganglia and midbrain (**Figure 1k-p**).

Neurons tended to be most active in a single position of the corridor and less active at the sensory identical position. Even when neurons responded in both repeating corridor segments, they often gave a larger response in one of the two segments than at the sensory identical position 40 cm away (e.g. **Figure 1e**). This preference was not explainable by random fluctuations, because the preferred positions used to sort neurons in the rasters (**Figure 1i-p**) were defined from held-out data (*24*, *25*). Thus, the presence of a strong main streak flanked by two weaker streaks 40 cm to the right or left confirms a reliable preference of the neurons for the first or second corridor segment.

This selectivity for a given position over the sensory identical position was present in all recorded regions. To measure selectivity, we centered the responses on each neuron’s preferred position, again defined from held-out data (*24*, *25*), and found that in each group of regions the average neuronal responses had a peak at the origin (the preferred position) surrounded by two smaller peaks 40 cm away (the sensory identical positions) (**Figure 1q-x, Figure S3, Figure S4**). To quantify this phenomenon, we calculated the Spatial Modulation Index (*24*, *25*): the difference between the responses at the preferred and sensory identical position, divided by the sum. Because the preferred position was computed from held-out data, this index has an expected value of zero for neurons with no reliable preference. However, as confirmed by a Linear Mixed Effects (LME) model, the spatial modulation index was significantly above zero in all recorded regions (LME with random effects of mouse and session: F_1,14767_ = 57.1, p = 4.4 x 10^- 14^) and was broadly similar across regions, ranging from 0.18 ± 0.01 in visual cortex (mean ± s.e., n = 1,802 neurons, with reliable responses across odd and even trials, **Figure 1r**) to 0.24 ± 0.01 in motor cortex (n = 2,148 neurons, **Figure 1t**). The hippocampal formation occupied a similar place as visual cortex (**Figure 1q**), with a spatial modulation index of 0.18 ± 0.01 (n = 5,043 neurons), which was significantly lower than in motor cortex (LME, F_36,14,767_ = 2.6, p = 3.7 x 10^-7^, post-hoc tests p < 0.01, **Figure 1y**). Its neurons thus did not stand out over the rest of the brain in their ability to correlate with spatial position beyond sensory inputs.

### Decoding position from any brain region

The appearance of a preferred position for neurons in all brain regions suggests that all these regions may carry a reliable correlate of position along the virtual corridor.

Consistent with this hypothesis, position along the corridor could be decoded from neural activity in virtually all recording sessions, regardless of the recorded regions. For each recording session, we trained a Bayesian decoder on the activity of all recorded neurons in a subset of trials and tested decoding performance on the held-out trials. For instance, in an example session that covered the dentate gyrus, CA3, and the dorsal lateral geniculate nucleus, the decoder correctly assigned high probabilities to the true position, with an overall accuracy of 43%, substantially above the chance level (9% for 11 bins, **Figure 2a**). Similarly, the decoding accuracy was 37% for another example session covering CA1, CA2, CA3, secondary motor cortex, somatosensory cortex (barrel field), and dorsal striatum (**Figure 2b**). On average, the decoding accuracy was 22.2 ± 0.6 % (mean ± s.e., 169 sessions) and was significantly above chance in 165 of 169 sessions (permutation test, p’s<0.05, **Figure S5a**). The four remaining sessions were excluded from other analyses.

**Figure 2.**
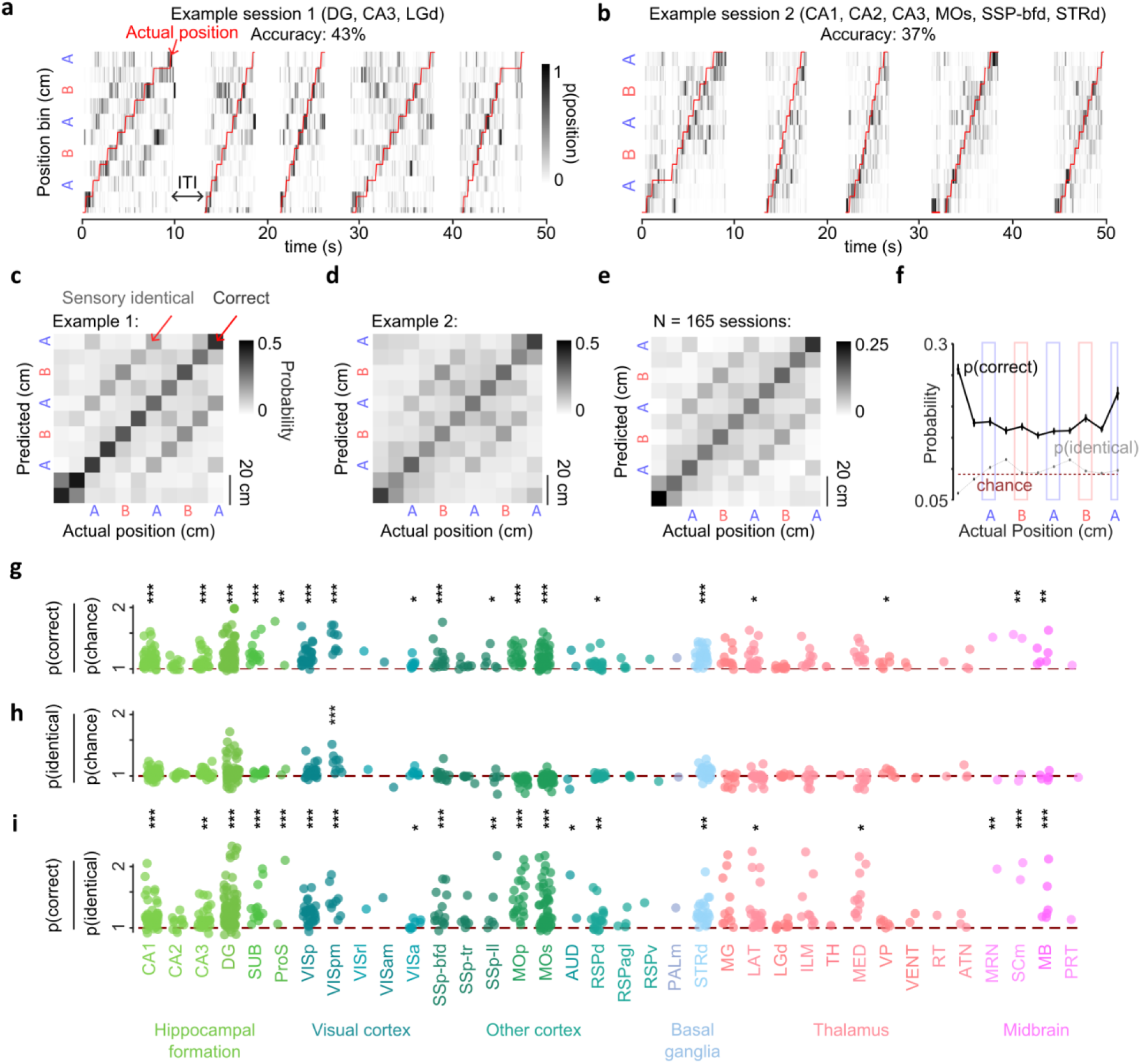
Decoding position from any brain region. **a)** Decoding probability as a function of position (ordinate, in 11 bins) and time (abscissa) for an example session with recordings in DG, CA3, and LGd. Actual position indicated by red line. Overall accuracy for session, 43%. **b)** Same, for another example session with recordings in CA1, CA2, CA3, MOs, SSP-bfd, and STRd. Overall accuracy 37%. **c)** Average probability of predicting a specific position given an actual position (confusion matrix) for example session 1. **d)** Same, for example session 2. **e)** Same, across 165 sessions. **f)** Probability of decoding the correct position (black) or the sensory identical position (gray), as a function of actual position (x-axis). **g)** Ratio between probability for correct position versus chance for individual sessions, grouped per brain region. **h)** Similar to g) but for the sensory identical position. **i)** Similar, but for the ratio between correct and sensory identical positions. Holm-corrected significance after establishing significant fixed effect of brain region in linear mixed effects model. *, p < 0.05; **, p<0.01; ***, p<0.001.

In all sessions, the decoder identified the correct position over the sensory identical position. In line with **Figure 1q-x**, the decoding probability was consistently higher for the correct position than for the sensory identical position (**Figure 2c-f**; post-hoc tests p < 0.05 after significant fixed effect in LME; F_2,492_ = 342, p = 7.4 x 10^-94^). As might be expected, nonetheless, the sensory identical positions were the most ambiguous: the decoded probability was higher at those positions than elsewhere (p < 0.05).

The accuracy of position decoding was above chance in every individual region, and significantly so in many cortical and subcortical regions. We trained decoders for each region separately, randomly subsampling equal numbers of neurons per region (**Figure S5b**). The decoding accuracy depended on the brain region (LME: F_35,422_ = 3.0, p = 6.9 x 10^-8^), and was above chance not only in the hippocampus, visual, somatosensory, dorsal retrosplenial, and motor cortices, but also in subcortical brain regions such as the striatum and some thalamic and midbrain regions (**Figure 2g**). Area VISpm, in particular, stood out as yielding significantly larger probabilities for the correct position than many other regions, including hippocampal regions (p’s < 0.05, post-hoc tests). This high performance partially reflected the encoding of visual inputs, since VISpm also yielded the highest probabilities for the sensory identical position (fixed effect of region, F_35,422_ = 4.7, p = 3.8 x 10^-15^, p’s < 0.05, post-hoc tests, **Figure 2h**). Regardless, across the brain the probabilities assigned to the correct position were larger than those to the sensory identical position. This was particularly the case for primary motor cortex (MOp), where (in line with the larger Spatial Modulation Index observed earlier, **Figure 1y**), the ratio between correct and sensory identical decoded position was larger than for hippocampal regions such as CA1, CA2, CA3, and DG (**Figure 2i**, fixed effect of area F_35,422_ = 2.4 p = 2.6 x 10^-5^, post-hoc tests p’s < 0.05).

### Visual anchoring of navigational signals

We next asked to what extent the position correlates across the brain were anchored to the sensory landmarks. An influence of the landmarks can be seen in the decoding accuracy, which was higher near landmarks than between them (**Figure 2f**). Between landmarks, the decoder assigned slightly higher probabilities to the sensory identical position. To investigate the impact of visual and auditory landmarks on the navigational signals, we randomly varied the visual contrast (0, 25, or 50%) and sound intensity (on or off) of the landmarks across trials.

Neuronal position correlates were strongly impacted by visual landmarks but scarcely by auditory landmarks. As illustrated by two example neurons in hippocampus (CA1) and visual cortex (VISpm), the visual contrast of the landmarks strongly modulated the responses, but the sound intensity had little effect (**Figure 3a,b**). This observation was consistent across neurons in those regions: position coding was invariant to the presence of the auditory landmarks, but it degraded as visual contrast was decreased, to the point that at zero contrast, position coding was only visible at the beginning of the corridor and near the end, after the reward position (**Figure 3c,d**). Accordingly, in the hippocampal formation and visual cortex, decoding accuracy increased with landmark visual contrast but did not vary with landmark sound intensity (**Figure S5c**). In neurons across the brain, average spatial tuning curves became substantially peakier at high landmark contrast but did not appreciably change with landmark sound intensity (**Figure S6**). The difference between the response at the preferred position and the average response increased with landmark visual contrast for all brain groups (LME models, as indicated in **Figure 3e**). However, it was unaffected by landmark sound intensity (p’s > 0.05, **Figure 3f**). A similar effect was observed for the difference in response at preferred versus sensory identical position (**Figure S7**). Similar results were seen at the level of individual brain regions (**Figure 3g**).

**Figure 3.**
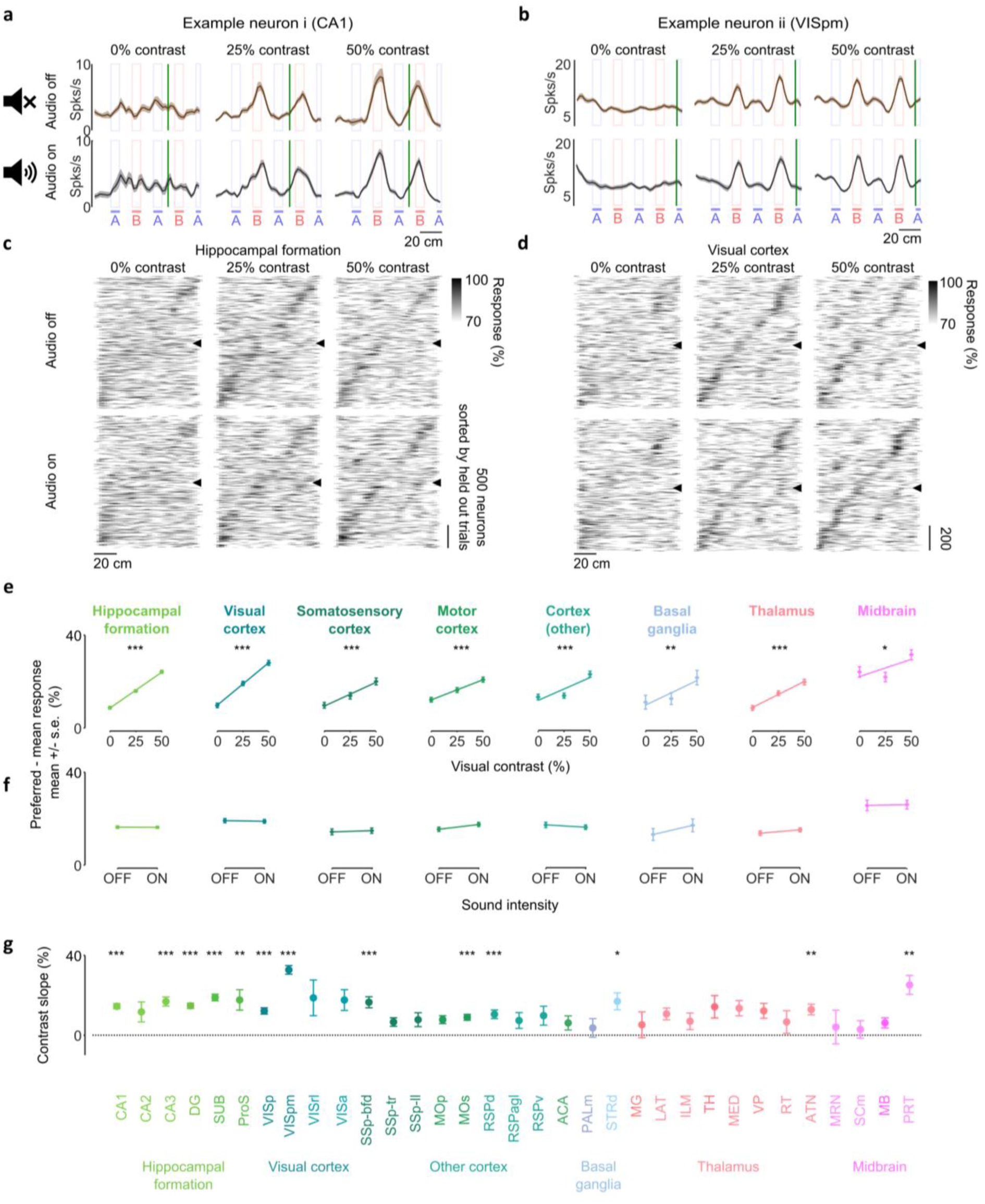
Visual anchoring of navigational signals. **a)** Response (mean ± s.e. across trials) as a function of position in the virtual corridor for example neuron in CA1 (same neuron as Figure 1e), shown for different landmark conditions in different panels. **b)** Same as a) for VISp (same as Figure 1f). **c)** Normalized activity of neurons in the hippocampal formation (y-axis) by spatial position (x-axis), separated for different trial conditions. Neurons were sorted by the position at which they maximally responded (i.e. preferred position) in held-out full contrast auditory-on trials. Only neurons with r>0 are included (see methods). **d)** Same as c) for neurons in the visual cortex. **e)** Difference (mean ± s.e.) between response at preferred position and mean response for visual contrast conditions, averaged across sound intensity conditions. Line is a linear fit to the data. ***, p<0.001; **, p<0.01; *, p<0.05 for significant effect of contrast (LME). **f)** Same as e) but for different sound intensity conditions, averaged across visual contrast conditions. **g)** Slope (mean ± s.e. across neurons) of the contrast fit as illustrated in e).

### Brainwide correlates of running

Having established that the brainwide correlates of position are strongly influenced by visual landmarks, we next asked whether they could be additionally explained by idiothetic signals such as the animal’s running speed. If running speed varies along the corridor, and if neurons correlate with running speed, these variations in speed might contribute to an apparent encoding of spatial position.

Mice did adjust their running speed along the corridor. Mice typically accelerated upon entering the corridor and decelerated as the end approached (**Figure 4a**). Moreover, the mice that received a reward (31 mice, 167 sessions) typically slowed down near the reward location (which differed across cohorts, **Figure 4a**). In 15 of the mice, this behavior was encouraged with double rewards (**Table S1**, see Methods). Indeed, running speed reflected an interaction between mouse position and reward position (mixed effects model, F_4,3662_ = 3.8, p = 0.004). This deceleration was largely due to expectations, as it was independent of actual reward delivery (not shown).

**Figure 4.**
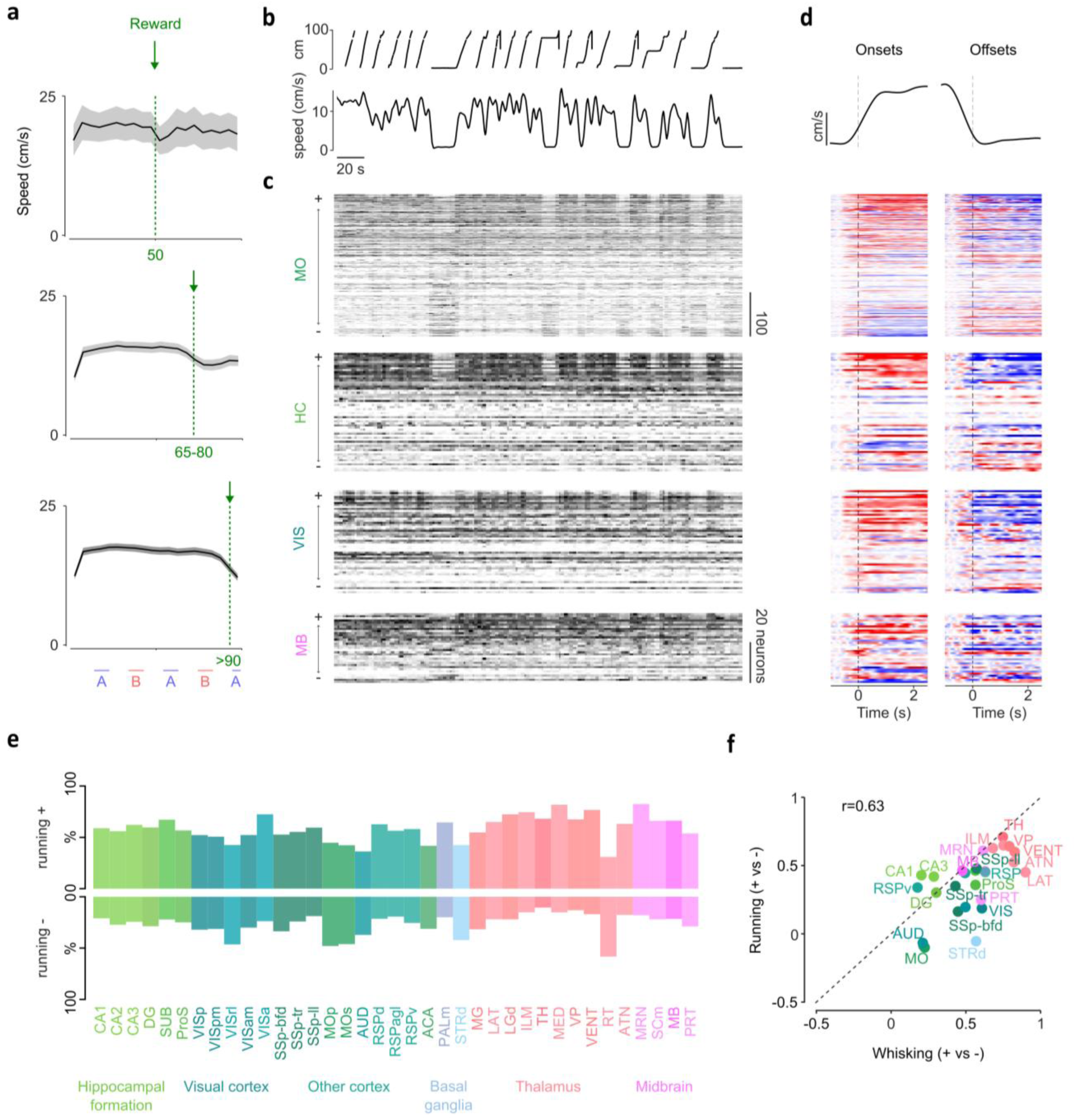
Brainwide correlates of running. **a)** Running speed as a function of position in the corridor, for sessions with different reward positions. **b)** Position (cm) in the virtual corridor (top) and running speed (bottom) over time (s) for example session. **c)** Same example session showing how individual neurons (rows) correlate their activity with running speed (b). Neurons are first sorted by brain region and then their correlation with running speed. **d)** z-scored activity aligned to onsets and offsets, sorted the same as in (c). Note that correlation was established in first half of the session, while activity aligned to on- and offsets was defined in second half of the session. **e)** Percentage of neurons positively (and significantly) correlated with running speed per brain region. **f)** The index between percentage of positive and negatively whisking-correlated neurons (data from(71)) against the same index for running-correlated neurons (new data).

Neurons in all regions showed positive or negative correlations with running speed. To observe these correlates of running consider an example session with recordings in motor cortex (MO), hippocampus (HC), visual cortex (VIS), and the midbrain (MB), where neurons are sorted within each region by their correlation with running speed (**Figure 4b,c**). In each region, a large fraction of neurons correlated positively with running, and another fraction, particularly visible in this session in MO, correlated negatively (**Figure 4d**). Such running correlations were widespread: every brain region contained neurons that correlated positively or negatively with running. The percentage of neurons with significant correlations (positive or negative) differed across regions (LME with binomial distribution, fixed effect of brain region: F_36,18806_ = 2.8, p = 7.0 x 10^-8^, **Figure 4e; Figure S8a,b**) and was highest in primary motor cortex (MOp). In most brain regions, more neurons correlated positively than negatively with running, particularly in CA3, lower limb primary somatosensory cortex (SSp-ll), and some thalamic regions (fixed effect of brain region: F_36,18806_ = 6.2, p = 7.2 x 10^-29^, and paired post-hoc tests). A possible exception was the reticular nucleus of the thalamus (RT), where more neurons correlated negatively than positively with running speed, but this difference was not significant (N=36 neurons, p > 0.05). In other brain regions such as the auditory cortex (Aud), motor cortex (MOp and MOs) and dorsal striatum (STRd), correlations with running speed were as often positive as negative (**Figure 4e; Figure S8a,b**).

These correlations of neuronal activity with running might be related to more general correlations with arousal levels. Neurons across the mouse brain correlate with arousal-related behaviors such as whisking, body or facial movements, even in mice that are not free to run (*51*, *52*, *69–71*). Intriguingly, the fraction of each region’s neurons that are positively vs. negatively correlated with running in our data correlated strongly with the fractions of neurons that are positively vs. negatively correlated with whisking in a different cohort of mice that were not allowed to run (*51*, *71*) (**Figure 4f; Figure S8c,d**). These similarities between correlates of running and whisking suggest that the brainwide correlates of running relate at least in part to arousal.

### Mixed encoding of sensation, reward, movement, and spatial position

We next sought to distinguish the encoding of spatial position from the contribution of vision, running, and other correlated factors. We fit the activity of every neuron as a weighted sum of factors that vary with time within each trial (*72*). These factors include spatial position along the corridor, sensory position (without distinction between the two identical segments), contrast and position of visual landmarks, the presence and position of auditory landmarks, corridor onset (which occurs in all trials), reward delivery (which is probabilistic but at the same positions across trials), distance run on the wheel (which depends on the random wheel gain), and running speed (**Figure 5a**). We fit the model with reduced-rank ridge regression (**Figure S9a**, see Methods), and measured the cross-validated correlation (cv-r) between the predicted and measured activity in held-out timepoints.

**Figure 5.**
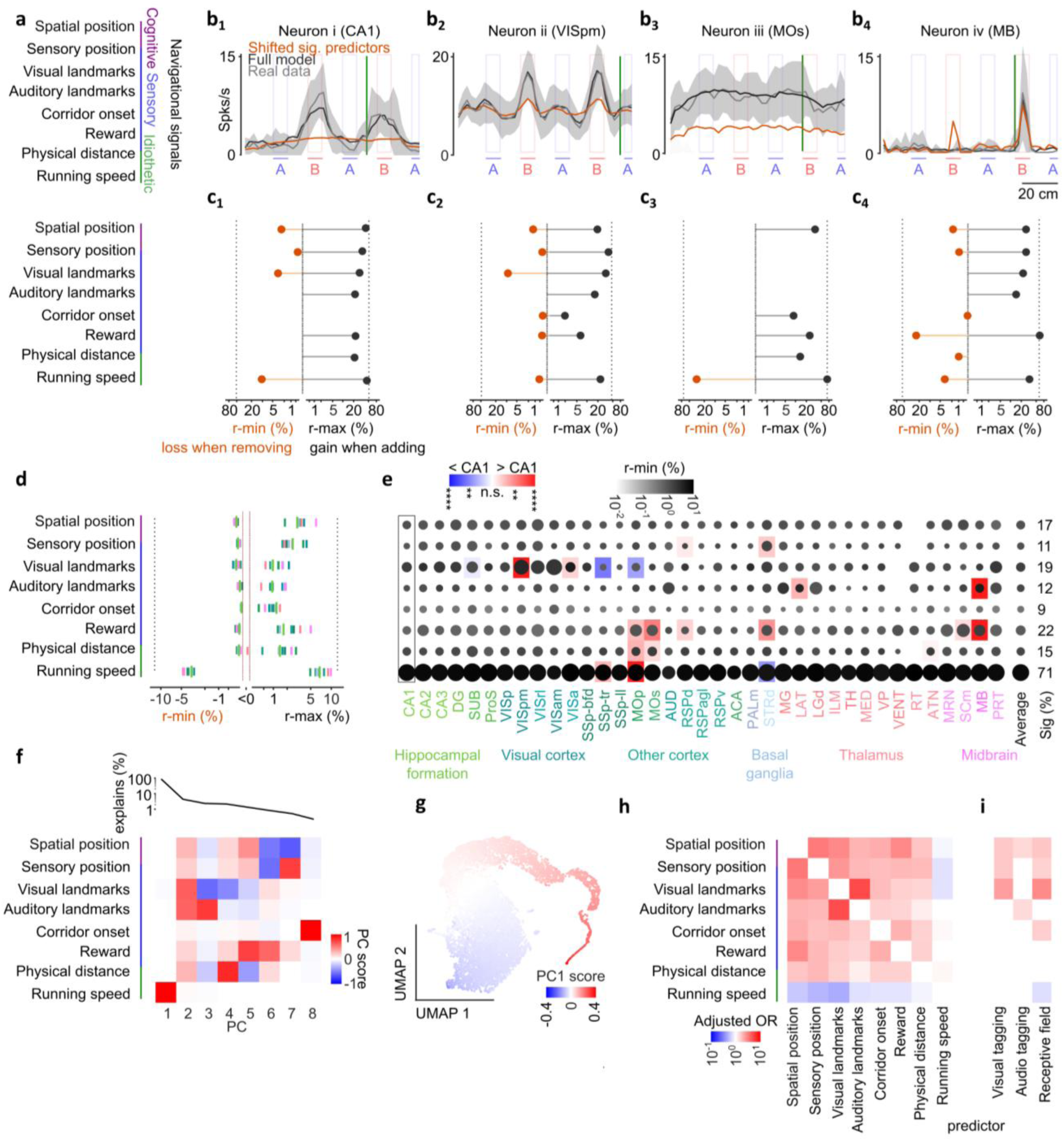
Mixed encoding of sensation, reward, movement, and spatial position. **a)** Predictor types in ridge regression model. **b)** Real data (gray, mean ± s.d. across full contrast no audio trials), full model prediction (black), model prediction with all significant predictors shifted (red, see c for which predictors these are). **c)** Loss (r-min) of removing a specific predictor from the full model (left axis), and gain (r-max) of having only the specific predictor in addition to the baseline firing rate. The cv-r of the full model is shown for comparison on both sides (dashed lines). **d)** Same as c) but as median per brain group. **e)** Bubble plot with relative size of the bubble indicating the percentage of neurons significantly encoding a variable (p < 0.05) and the color (light to dark gray) indicating fixed-effect estimates of r-min. Areas with significantly different r-min values from CA1 are indicated by a red-blue square background, p-values from post-hoc tests after LME. For comparison the average percentage of neurons significantly encoding each variable across brain regions is also indicated. **f)** PCA-analysis performed on r-min profiles. Top: explained variance of each PC. Bottom: score per predictor per PC. **g)** 2D-UMAP representations (minimal distance: 0.5, number of neighbors: 5) of r-min profiles, each data point is a neuron colored by the score of PC1, which was fully loading on running speed (see f). **h)** The adjusted odds ratio (mixed-effects model estimate, see Methods) of a neuron being already selective to one navigational variable (x-axis) to be co- selective for another one (y-axis). **i)** Adjusted odds ratio for neurons significantly tuned to passive stimuli, to also encode any of the navigational signals. Only odds significantly different from one are shown. See **Figure S10** for details about passive stimuli and example responses.

This model revealed that most neurons across the brain correlated with at least some of the navigational signals. The model’s performance can be observed in the four example neurons recorded in CA1, VISpm, MOs, and MB, where the model provided cross-validated correlations (cv-r) between 50% and 70% (**Figure 5b**). Model fits were particularly good in subiculum (SUB), VISpm, MOp, anterior group of dorsal thalamus (ATN) and midbrain reticular nucleus (MRN) (p < 0.05 after main effect; F_36,18806_ = 9.1, p = 4.8 x 10^-48^, **Figure S9b**). The cv-r was positive in 90.2% of all neurons, indicating that these neurons were modulated by one or more navigational signals. The remaining 9.8% of the neurons were presumably unrelated to navigation and were excluded from further analysis.

We then tested the role of the individual factors by removing them from the full model and by considering them in isolation (*70*, *72*). First, to estimate whether a factor made a unique contribution to neuronal activity beyond other correlated factors, we circularly shifted that factor before fitting the model. The difference in performance (cv-r) relative to the original fit establishes that factor’s minimal contribution (r-min). Second, to estimate the potential contribution of a factor in isolation, we considered a reduced model that only had that factor plus a constant. The performance (cv-r) of this reduced model establishes that factor’s maximal contribution (r-max). As expected, this second measure was more generous than the first. For instance, for example neuron *i*, the maximal contribution was sizeable for all factors except for corridor onset, but the minimal contribution was sizeable for only four factors: spatial position, sensory position, visual landmarks, and running speed (**Figure 5c_1_**). Indeed, an additional fit where we shifted all those four factors led to visibly worse predictions (**Figure 5b_1_**). For example neuron *iii*, the minimal contribution was sizeable for only one factor: running speed (**Figure 5c_3_**), and indeed, shifting that factor visibly worsened the predictions (**Figure 5b_3_**).

Running speed was the best individual brainwide predictor of neural activity, but it was unable to explain the responses without other navigational signals. In line with the brainwide correlates of running that we had observed (**Figure 4**), running speed explained most variance in neural activity, providing a sizeable median contribution to both r-max (9.8 ± 2.1%; median ± m.a.d. across 37 regions) and r-min (6.3 ± 0.9%) across the brain (**Figure 5d**). The minimal contribution r-min of running speed was largest in primary motor cortex (MOp) and smallest in the dorsal striatum (STRd), with CA1 in an intermediate position (**Figure 5e**). Nonetheless, only in one region (ventral posterior thalamus, VP) was it able on its own to explain neural activity as well as the full model (LME, interaction of brain region x model type: F_36,37612_ = 18.0, p =1.7 x 10^-112^, post-hoc tests p < 0.05, **Figure S9b**). In all other brain regions, the responses were also driven by other navigational signals.

Additional navigational signals that were necessary to explain brainwide activity included reward, visual landmarks, and spatial position. The contributions to r-min and r-max of other navigational signals were typically lower than those of running speed, with median r-max at 1-5% (**Figure 5d**). The median r-min was much lower, below 0.5%, indicating that these navigational signals are correlated, with r-max overestimating and r-min underestimating their contribution. To be conservative, we defined a neuron’s ‘encoding’ of a factor based solely on a significantly positive r-min. With this criterion, running speed was encoded in 71 ± 2% (mean ± s.e. across regions) of neurons with the largest r-min (6.3 ± 0.9%) and r-max (9.8 ± 2.1%). Reward was encoded in 22 ± 1% of neurons, with minimum contributions of 0.3 ± 0.1% and maximum contributions of 3.8 ± 0.9%, and was a particularly strong predictor in midbrain regions, striatum, and motor cortex. In combination, therefore, running speed and reward explained the strong spatial modulation (**Figure 1y**) and decoding accuracy (**Figure 2i**) of motor cortex and midbrain. Visual landmarks were encoded in 19 ± 2% of neurons, with minimum contributions of 0.5 ± 0.2% and maximum contributions of 2.4 ± 1.1%. As expected, the visual cortex had the strongest visual contrast encoding, higher than the hippocampal formation (**Figure 5e**). Conversely, SSP-tr and MO had significantly weaker visual contrast encoding than the hippocampal formation. Spatial position, on the other hand, explained activity more equally across the brain: in 17 ± 1% of neurons, with unique contributions r-min of 0.4 ± 0.1% and maximum contributions r-max of 5.1 ± 1.3% (**Figure 5e**).

Neurons across the brain showed mixed selectivity for multiple navigational signals, with a principal axis of variation determined largely by running speed. In most neurons the activity correlated with more than one navigational signal (**Figure S9c**). Principal component analysis revealed that running speed explained most of the variance in the r-min values, thus defining the main axis of variation in tuning profiles across neurons (**Figure 5f**). This axis of variation was evident in a low-dimensional representation of the neuronal population (UMAP, **Figure 5g**). Running speed was also the signal that was most separable: a neuron selective for running speed was not more likely to be selective to any other navigational signal (**Figure 5h**). For all other navigational signals, neurons were typically co-tuned: neurons encoding spatial position were more likely to encode sensory position, visual landmarks, and rewards (**Figure 5h**, **Figure S9d-e**).

Neurons with passive visual tuning were more likely to encode sensory position and spatial position. We presented passive stimuli to the mice after some of the sessions in the navigation task, including receptive field mapping and the sensory stimuli used in the virtual corridor. Sensory stimuli were presented at a specific frequency to allow frequency tagging (*73*): a neural response phase-locked to the specific frequency can be attributed to that specific sensory input. Neurons responding to these passive stimuli (e.g. **Figure S10a-o**) were more prevalent in some regions than in others. Visual frequency tagging had particularly high prevalence in the hippocampal formation, visual cortex, and basal ganglia (χ^2^(7) = 520.6, p < 0.001, corrected post-hoc tests, **Figure S10p**), and low prevalence in the motor cortex. Neurons significantly tagged by visual stimuli were more likely to encode visual landmarks, sensory position, and spatial position in the corridor. Neurons significantly tagged by auditory stimuli also encoded auditory landmarks and position in the corridor. The number of neurons with receptive fields was significantly larger in visual cortex and the hippocampus (χ^2^(7) = 180.6, p < 0.001, **Figure S10r**). Neurons with receptive fields were more likely to be tuned to visual landmarks, sensory position, and spatial position in the corridor (**Figure 5i**). These findings suggest that neurons with visual responses are more likely to be recruited in navigation.

Thus, in addition to a brainwide encoding of running, there is a brainwide encoding of position. Position encoding is more likely in neurons that encode sensory stimuli, even outside of a task.

### Uniformity and precision of spatial representations

Having found that a similar percentage of neurons encode position brainwide, we next asked whether some regions represent the environment better than in others. We defined a better representation as one where the preferred positions of the neuronal population provide a more uniform coverage of the corridor (*74*), and one where the population vector encodes position more precisely (*75*, *76*).

The most uniform coverage of the corridor was in the hippocampal formation. In all 8 groups of regions we observed some clustering of preferred response positions near the corridor onset, perhaps reflecting a general change in brain state associated with the audiovisual stimulation (*77*) (**Figure 6a**). For the following analyses we therefore focused on positions >5 cm into the corridor. The distributions of preferred positions tended to cluster near the landmarks (**Figure 6a**) and near the reward position (**Figure 6b**). Permutation tests showed that alignment to the nearest landmark was prominent in visual cortex and the basal ganglia (permutation test, p < 0.001), and low for somatosensory, motor, and other cortex (p < 0.05, **Figure S11a**). Alignment to reward was prominent in somatosensory cortex, other cortex, basal ganglia, thalamus and the midbrain (permutation test, p < 0.001), and less in motor cortex and hippocampal formation (p < 0.01, **Figure S11b**). To quantify uniformity, we computed the normalized entropy of 200 randomly sampled neurons and expressed it as a percentage of maximally expected entropy, repeating this procedure 50 times (*74*). The distribution of preferred response positions was more uniform in some brain regions than others (χ^2^(7) = 187, p = 0.001, permutation test of the group effect). Coverage was most uniform in the hippocampal formation (97.2 ± 0.15%), with motor cortex (96.8 ± 0.14%) close behind. Regressing out running speed left these results mostly unchanged (χ²(7) = 148, p = 0.002), leaving the hippocampal formation in the lead (97.0 ± 0.18%) but also revealing a similarly uniform coverage in the thalamus (97.2 ± 0.14%).

**Figure 6.**
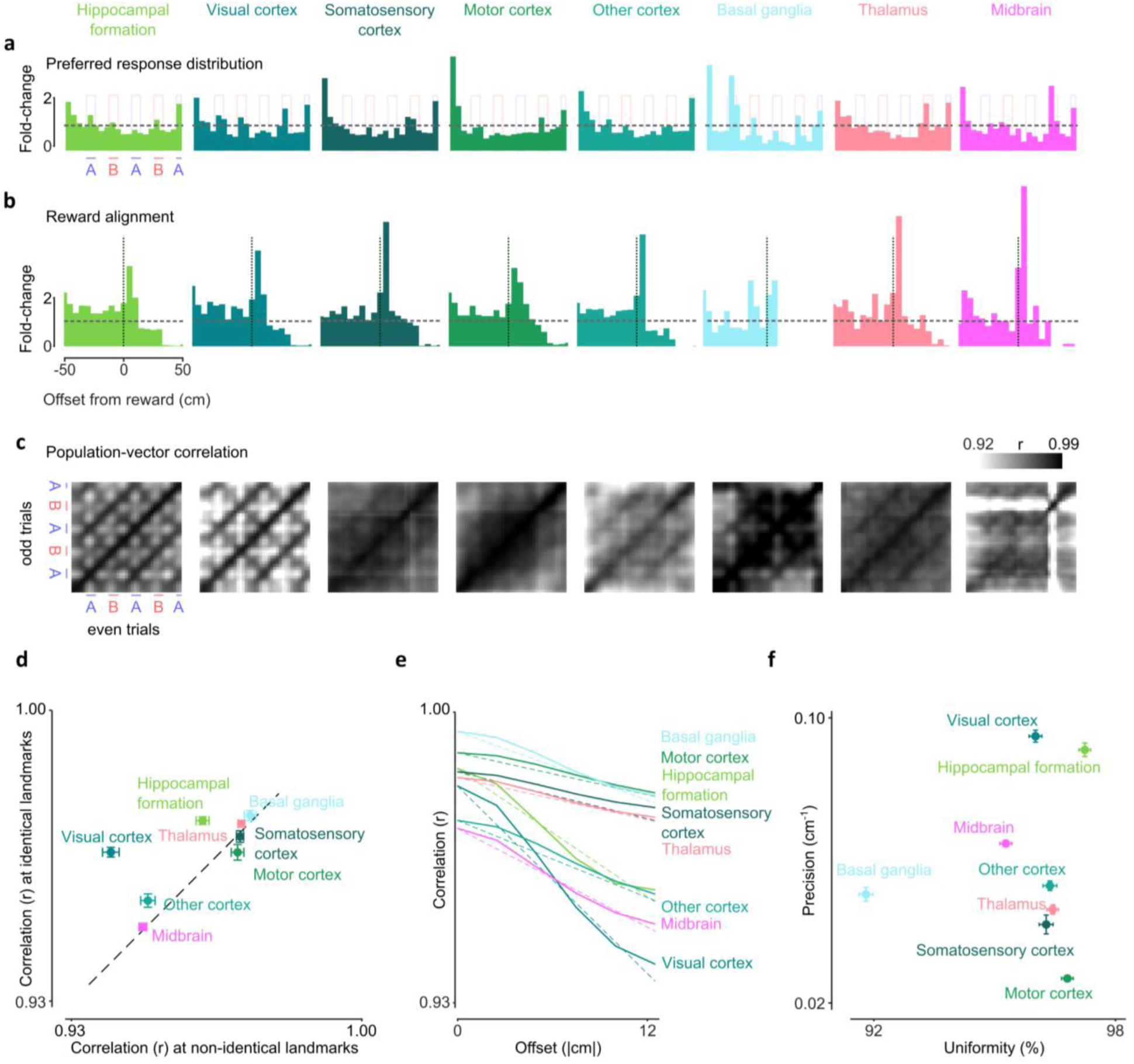
Uniformity and precision of spatial representations. **a)** Histogram of preferred response positions (fold-change relative to expected from a uniform distribution, indicated by the dashed horizontal line). **b)** Histogram of preferred response positions aligned to reward delivery. c) Population-vector correlation of mean activity between odd and even trial responses of 200 randomly sampled neurons x 39 position bins at 2.5 cm resolution for every pair of positions. See Figure S11c for individual brain regions. **d)** Population-vector correlation for sensory-identical (offsets at 40 and 80 cm) positions versus non-identical (offsets at 20 and 60 cm) positions. **e)** Correlation as a function of spatial offset (|cm|) (solid), and a linear fit to estimate the spatial precision (i.e. slope of the decline in cm-1). **f)** Average precision (mean ± s.e. across draws) as indicated in e) against uniformity (%), defined as normalized entropy where 100% is the maximally expected entropy. Higher values indicate more uniformity/precision of responses, lower values more clustering/less precision. See Figure S11d for individual brain regions.

We then measured the variation of the population responses along the corridor and found it to be highest in visual cortex and hippocampal formation. To assess the similarity of population responses across position, we pooled neurons across sessions, drew matched-size random subsets per brain group (N = 200, 50 draws), and built spatial tuning curves separately from odd and even trials. We then computed the population-vector correlation between odd- and even-trial tuning curves at each pair of positions (*75*, *76*), thus obtaining a matrix of correlations where high values indicate similar population responses (**Figure 6c**). The results revealed sharp bands with high correlations at corresponding positions (the diagonal), indicating that the population response at any given position was highly reliable. Reliability was particularly high (0.99 ± 0.001) in hippocampal formation, motor cortex, somatosensory cortex, and basal ganglia. Similar results were seen in individual regions (**Figure S11c**). The population responses were also quite similar at sensory identical positions (off-diagonal bands), especially in the hippocampal formation and in the visual cortex, and, to some extent, basal ganglia and other cortex (**Figure 6c,d**).

In visual cortex, hippocampal formation, and basal ganglia, the population responses were specifically driven by the visual landmarks. Indeed, in these groups of regions the population responses at the landmark positions tended to differ from the responses at other (sensory different) positions (lighter horizontal and vertical bands, forming crosses, **Figure 6c,d**). By contrast, in midbrain the population responses differed from all other positions only at the reward position. The high landmark specificity of the population responses in visual cortex, hippocampal formation, and basal ganglia indicates that these regions are specifically driven by landmarks. Given the previous findings (**Figure 3**, **Figure 5**), most likely the visual contrast of these landmarks is what drove the high specificity rather than the sound intensity. Indeed, these regions had a particularly large fraction of neurons that were selective for visual landmarks in the passive condition (**Figure S10p**).

The visual cortex represented spatial position more precisely than the hippocampal formation. We measured each region’s spatial precision as the rate at which the population-vector correlation (min-max normalized) decayed with spatial offset (**Figure 6e**). Spatial precision differed significantly across the eight groups of regions (χ²(7) = 306, p = 0.001). Visual cortex had the highest precision (0.095 ± 0.002 cm^-1^), followed closely by the hippocampal formation (0.091 ± 0.002 cm^-1^). The somatosensory cortex (0.042 ± 0.003 cm^-1^) and motor cortex (0.027 ± 0.001 cm^-1^) had the least precise representation (**Figure 6f**). Across all individual regions, the highest precision and the highest uniformity was in dentate gyrus (DG), closely followed by CA1 and primary visual cortex (VISp) (**Figure S11d**). These results were robust to correcting for running speed: visual cortex and hippocampal formation remained the two most precise regions (χ²(7) = 297, p = 0.001).

In summary, while the hippocampal formation was far from having the highest percentage of neurons encoding spatial position, it had the most uniform coverage of the corridor, and its spatial code was the second-most precise.

## Discussion

By recording from thousands of neurons across the mouse brain while decoupling position from associated signals in a virtual corridor, we revealed a highly distributed organization of navigation signals. Neurons in all brain regions gave stronger responses in one position than in others, typically tiling the corridor. This selectivity for position was reliable, allowing the mouse’s position to be decoded from the activity of any region. It was anchored by the onset of the corridor, by the visual (but not auditory) landmarks, and by the reward position. It was also powerfully shaped by running, which correlated positively or negatively with neurons in all regions and was the strongest determinant of responses among the navigational signals. The resulting brainwide representation of navigational signals was largely composed of mixed-selective neurons tuned to combinations of running, sensation, reward, and position. Vision played a particularly strong role, as selectivity for position during navigation correlated with visual responses measured in the absence of navigation.

A simple additive model revealed that spatial position is represented in every region, and that its representation is mixed with other factors. Mixed selectivity has emerged as a general feature of cortical computation in tasks involving multiple variables (*78*, *79*) and has potential functional advantages: it increases the dimensionality of population codes and allows downstream readouts to flexibly combine variables. Mixed selectivity for navigation signals had long been observed in the hippocampus where neurons encode not only spatial position but also non-spatial signals (*36–38*, *80–82*). Here we discovered that this mixed selectivity for spatial and non-spatial signals occurs brainwide.

Among the other factors contributing to brainwide activity during navigation, the strongest one – and the least mixed with other factors – was running speed, which correlated positively or negatively with the activity of almost all neurons across the brain. This result extends previous observations of running modulation in primary visual cortex (*83*) and in other cortical areas (*53*, *54*) including entorhinal cortex (*84*, *85*). These widespread correlates of running echo the correlates of whisking and other uninstructed movements (*69–71*, *86*), and of task movements (*52*), which are also brainwide. The number of neurons that correlated positively vs. negatively with running differed across areas and was approximately equal in auditory cortex (Aud), motor and prefrontal cortex (MOp and MOs), and dorsal striatum (STRd). We found it to be strongly related to the number of neurons that correlate positively vs. negatively with whisking in stationary mice (*51*, *71*), suggesting that both effects reflect arousal rather than specific motor signals. Indeed, consistent with previous observations that the correlates of arousal are mostly orthogonal to task factors (*69*), we found that the correlates of running were largely orthogonal to the other factors.

Another key factor that determined brainwide position signals were the visual – but not auditory – landmarks. The visual landmarks strongly impacted the neuronal responses. When we decreased their contrast, we observed a marked degradation of position coding across the brain, to the point that at zero contrast, position coding was only visible at the beginning of the corridor and after the reward. Vision thus informed a representation of spatial position that was distributed brainwide.

Conversely, position coding (and decoding) was scarcely if at all affected by the auditory landmarks. We found this surprising, given that many of the same neurons across the brain responded in a frequency-dependent way to the auditory stimuli used in the virtual corridor, and that neurons in the hippocampus can follow auditory stimuli (*50*, *87*). Perhaps in our corridor the auditory landmarks were not particularly helpful, because their superimposed sounds were audible from the entire corridor, varying only in relative amplitude in the different positions. It is possible that if the auditory landmarks had been fewer or spaced further apart, the neurons might have relied on them more.

Vision thus informed a representation of spatial position that was distributed brainwide. Just as we might consult a map for different reasons in different contexts, a distributed representation of position across the brain could be exploited in different ways depending on the region and the context, to locate diverse features such as rewards (*38*, *39*) or objects (*6*, *88*). In the context of our experiment, a linear virtual corridor, the hippocampal formation’s population tuning was nearly as precise as that of the visual cortex, and many neurons in these regions were tuned to visual stimuli both during navigation and in the absence of navigation. Other regions clustered their responses more around reward locations than landmarks, but without the presence of visual landmarks, spatial preferences still collapsed for most of these regions.

Given this brainwide representation of position, does the hippocampal formation play a special role in navigation? If our mice had been in a real-world 2D environment, we would have been able to measure additional navigational signals such as those related to head direction, which are much stronger in some regions than others (*89–91*). In 2D, the nature of spatial encoding in regions of the hippocampal formation such as entorhinal cortex would likely surpass that of many other regions. But in our 1D virtual track, it hardly did. Indeed, the hippocampal formation did not contain more neurons encoding position than other regions. Moreover, the activity of its neurons was not explained by position better than in other regions. As a population, however, its neurons displayed a more uniform representation of the corridor, with the least tendency to cluster preferred responses to specific reward locations and landmarks.

Finally, the hippocampal formation’s tuning was very precise. It is thus possible that the hippocampus is itself the source of the spatial representation present in all other brain regions (*14*, *92*). Our results, however, reveal that it is far from unique in containing a spatial representation: spatial position and other navigational signals are represented extremely widely across the brain.

## Acknowledgements

We thank Ran Bi for assistance in some of the recordings, Charu Bai Reddy for lab management, Michael Krumin for technical help, BSU staff for animal welfare monitoring and care, and members of the Carandini-Harris and Barry labs at UCL for useful discussions and feedback. Claude, Codex and Gemini assisted the authors with analyses and a few textual suggestions. This research was funded by the Wellcome Trust (Investigator Award 223144/Z/21/Z to K.D.H. and M.C.) and the European Union’s Horizon 2020 (Marie Skłodowska-Curie grant agreement no. 101022757 to E.H.vB.). M.C. holds the GlaxoSmithKline / Fight for Sight Chair in Visual Neuroscience.

## CRediT contributions

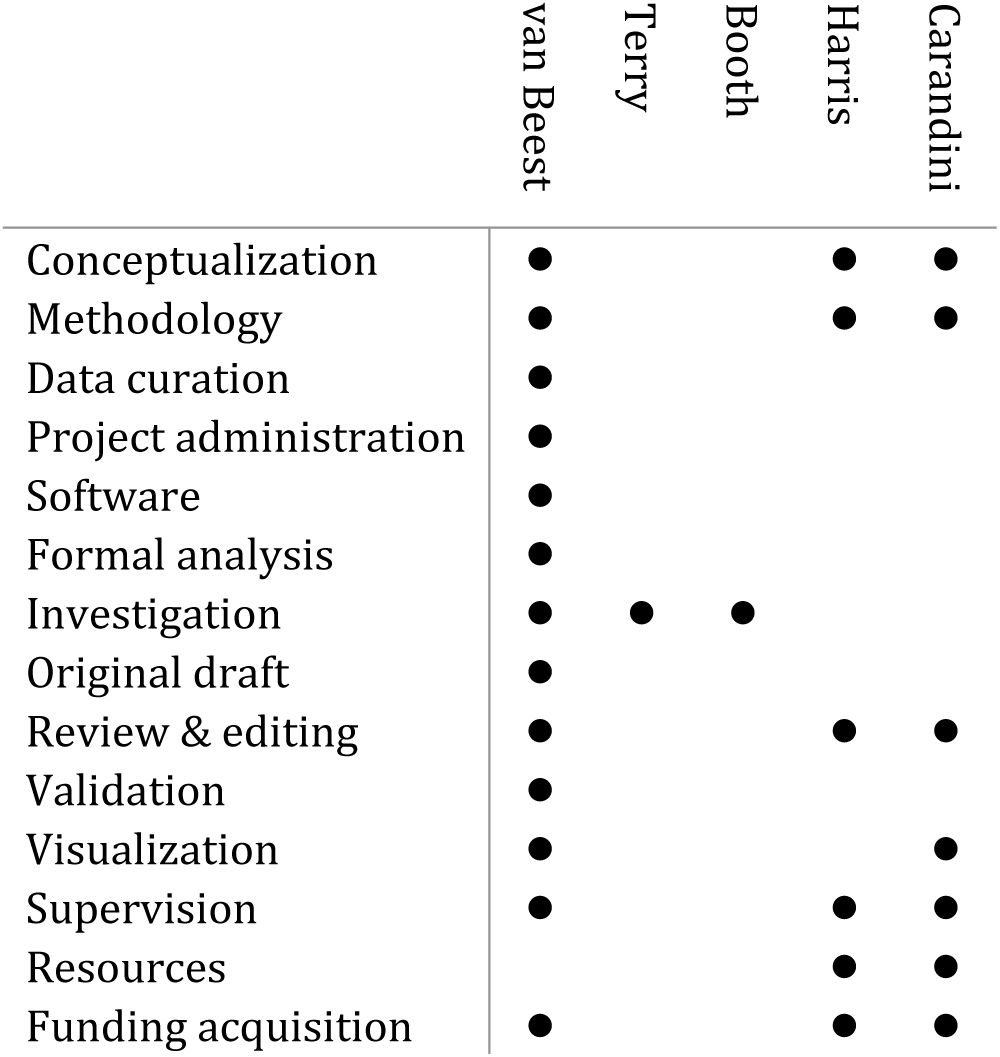

## Data, code, and materials availability

Spike-sorted data and code to reproduce results will be released upon publication.

