## Supplementary Materials for "Brainwide representation of navigation"

Materials and Methods

Figs. S1 to S12

Tables S1 to S2

References (93 – 103)

#### Materials and Methods

Experimental procedures were conducted at UCL according to the UK Animals Scientific Procedures Act (1986) and under licenses released by the Home Office following appropriate ethics review. We used 28 C57BL/6 mice (12 female, **Table S1**) aged 1.4 to 7.2 months at the time of first surgical procedure. Of these, 17 were recorded acutely (a probe was inserted and retracted for every recording session), and 11 chronically (a probe was inserted stably for months) (*95*). Seven of the mice also participated in other studies (not presented here).

##### *Surgeries*

A brief (~ 1 h) initial surgery was performed under isoflurane (1–3% in $O_2$) anesthesia to implant a titanium headplate (~ 25 × 3 × 0.5 mm, 0.2 g for chronic implants or well-shaped ~14 x 5-14mm, 0.9 g for acute implants). The surface of the skull was cleared of skin and periosteum and covered with a layer of cyanoacrylate followed by layers of UV-curing optical glue (Norland Optical Adhesives #81, Norland Products). The head plate was then attached over the interparietal bone with Super-Bond polymer.

*Chronic implant*

Chronic implants were performed with a modular recoverable implant (*95*). Prior to surgery, the probes were coated with DiI by brushing them with a droplet of DiI or dipping them in DiI, for histological reconstruction. Craniotomies were performed on the day of the implantation or one day before, under isoflurane (1–3% in O2) anesthesia, and after injection of Colvasone and Rimadyl. The UV-curing glue was removed, and the skull cleaned and scarred for best adhesion of the cement and levelled. Craniotomies were performed with a drill or a biopsy punch, and the exposed brain was covered with Dura-Gel (Cambridge Neurotech). The implant was held using the 3D-printed payload holder (*95*) and positioned using a micromanipulator (Sensapex). The probes were positioned at the brain surface to avoid blood vessels and then inserted slowly (3-5 μm/s). Once the desired depth was reached (ideally just before the docking module touched the skull), the implant was sealed using UV glue and covered with Super-Bond polymer, ensuring that only the docking module was cemented. After finishing all recording sessions, the probes were explanted and cleaned before reusing.

*Acute preparation*

Craniotomies as described above were performed at least four hours prior to the first recording, and typically >12 h before, and sealed first with Dura-Gel (Cambridge Neurotech). KwikCast was applied and a plastic lid were screwed on the headplate in between recordings. Prior to probe insertion, the probes were coated with DiI by brushing them with a droplet of DiI or dipping them in DiI, for histological reconstruction.

After recovery of every surgical procedure, mice were treated with carprofen or meloxicam for three days, then acclimated to handling and head-fixation.

#### ***Recordings***

##### *Experimental rigs*

During the recording sessions, mice were head-restrained in the center of three 10-inch LCD screens (1024 x 768 pixels, Adafruit, LP097QX1) placed at 90° angles. The distance from each screen was 11 cm, so that the screens covered the visual field by 135° in azimuth and 45° in elevation. The screens were fitted with Fresnel lenses (Wuxi Bohai Optics, BHPA220-2-5) and a polarizing filter (Rosco) to provide approximately equal intensity across viewing angles. The refresh rate was 30 Hz in virtual reality (VR) and 60 Hz in passive stimulus presentations. Intensity values were linearized with a photodiode (PDA25K2, Thorlabs).

Auditory stimuli were presented by a speaker (102-1299-ND, DigiKey) positioned below the screens in front of the animal (0° azimuth). Its frequency response was estimated with a calibrated microphone (GRAS 40BF ¼" Ext. Polarized Free-field microphone).

##### *Virtual reality*

The VR corridor contained two sensory identical segments (*24*). The corridor was 8 cm wide and 100 cm long. The walls of the entire corridor were covered with a background Gaussian noise texture (50% contrast) which was always present and which added to the visual landmarks when these were present. It started with 16 cm of open corridor (i.e. with no landmarks), followed by a 40 cm segment AB, by another identical segment AB, and by 4 more identical segments AB to ensure identical audiovisual experiences in the first two segments. The corridor thus appeared to be 256 cm long at the onset, but actually ended after 100 cm. Each segment AB contained an 8 cm section with landmark A, a 12 cm open corridor, an 8 cm section with landmark B, and a 12 cm open corridor. Landmarks A and B were thus 20 cm apart, and the complete segment AB was 40 cm long. The segment was repeated at 16, 56, 96, etc. cm from the start of the corridor, and the 100 cm traversable corridor contained landmark A centered at 20, 60, and 100 cm and landmark B centered at 40 and 80 cm.

The landmarks A and B were both visual and auditory. The visual landmarks were a grating (A) and a plaid (B). In some sessions, these landmarks flickered at a frequency of 3.35 (A) or 6.11 (B) Hz (*73*). In each trial, the contrast of all the visual landmarks was set to 0, 25, or 50%. The auditory landmarks were an 11 kHz tone cosine-modulated at 5 Hz (A) and a 17 kHz tone cosine-modulated at 7.07 Hz (B). Sound intensity scaled with distance to the source location (e.g. A or B) according to the inverse-square law (default setting in the openAL integration of Psychtoolbox-3), to maximally 65 dbSPL at the center of A/B landmarks. Just as the visual landmarks were visible far into the virtual corridor, all auditory landmarks were audible from the onset of the corridor. In each trial, the auditory landmarks could be present or absent. To pilot a different study, in some sessions, a small subset of the trials included 3D objects placed 8 cm before the original landmarks (a teapot near the left wall (A) or a cube (B) near the right wall). Data from those trials does not pertain to this study.

Animals traversed the virtual corridor by walking on a transparent wheel (19 cm diameter, (**https://hackaday.io/project/160744-kinemouse-wheel**) or an anodized aluminum mesh wheel (20 cm diameter, designed by Charu Bai Reddy and Pip Coen). Running speed was measured online with a rotary encoder (50 or 1024 rps, Kübler, Germany) to update the virtual scene, with a variable gain randomly sampled between 0.5:1 and 2:1. Upon reaching 100 cm into the corridor, the screens turned gray for a random inter-trial interval of 2-3 s, after which the mice were placed at the beginning of the virtual corridor for the next trial. The duration of each trial depended on how long it took the mouse to traverse the corridor. In some sessions, if the end of the corridor was not reached within 30 s, the corridor passively moved forward with a speed between 5 and 10 cm/s (0.2 cm/s spacing, randomly sampled). In other sessions a time-out occurred after 30 or 60 s. A typical session included 50-250 trials.

To motivate mice – which were water limited - to run, a water reward was delivered in the corridor, typically between 70 and 95 cm (for a given session always in the same location) with a probability of 0.85-0.90. Other than reaching the reward location, no action was required to obtain the reward. In some sessions, mice received double the reward if they slowed down at its location (**Table S1**).

Mice were handled and accustomed to the experimental rigs prior to experiments. Acutely implanted mice were familiarized with the virtual corridor prior to recordings. Chronically implanted mice were familiarized with running in a corridor with the same noise background but without landmarks to obtain rewards prior to implantation, and then with the full virtual corridor after implantation. The recordings analyzed here were obtained in mice that were familiar with the corridor.

##### *Passive stimulation*

To establish tuning properties of neurons outside of the virtual corridor, after some of the sessions of virtual navigation we presented passive stimuli. The passive stimuli included receptive field mapping and frequency tagging of the auditory and visual landmarks used in the virtual corridor (*73*).

Receptive fields were mapped using a sparse-noise stimulus: a binary checkerboard grid refreshing at 60 Hz, with each frame independently and randomly assigning each spatial position (5 deg squares) as ON (white), OFF (black), or blank (gray). Stimulus frames were reconstructed offline by detecting photodiode flip times and assigning each inter-flip interval to a stored frame index. For each neuron, the mean spike count at each grid position was computed separately for ON and OFF frames within a response window following stimulus onset, yielding raw ON and OFF receptive field maps. Each map was z-scored across all spatial positions. A combined map was derived as the sum of the absolute z-scored ON and OFF maps, then z-scored again. All maps were spatially smoothed with a 2-D Gaussian kernel (smooth2a). A rotated 2-D Gaussian was fit to each map using nonlinear least squares (lsqcurvefit with D2GaussFunctionRot), estimating six parameters: amplitude, center (azimuth, elevation), tuning widths, and rotation angle. Statistical significance was assessed by bootstrap: spike-time labels were randomly shuffled, RF maps were recomputed and fit for each shuffle, and the residual norm of the actual fit was compared against the shuffle distribution to derive a p-value.

To establish whether the sensory stimuli used in the VR elicit significant responses in recorded neurons, we presented random combinations of the visual and auditory landmarks used in the virtual corridor as described above, except visual stimuli were presented full-screen, 0.12 cpd, at 80% contrast. All stimuli were presented at their assigned frequency for frequency tagging (*73*), for a total duration of 2 s. To test for significant frequency tuning, we extracted the phase of all spikes relative to the frequency of the stimulus and used the Rayleigh test for non-uniformity of circular data and the circular mean computation from the Circular Statistics toolbox for MATLAB (*96*).

#### *Data processing*

Electrophysiology data were acquired using SpikeGLX (`https://billkarsh.github.io/SpikeGLX/`) and spike sorted with Kilosort4 (*97*). Data were preprocessed using the default pipeline available via UnitMatch (*68*) to merge oversplits (`https://github.com/EnnyvanBeest/UnitMatch`), and the selection of well-isolated single neurons with Bombcell (`https://github.com/Julie-Fabre/bombcell`)(*67*, *68*). All data including spike times were down sampled to 30 Hz. Only sessions with at least 25 trials that reached at least 85 cm of the virtual corridor were included. Due to the nature of chronic implants, we sometimes found neurons that were present in multiple sessions (*68*). We retained one representative session per neuron to avoid pseudo replication in later mixed-effects models: among a neuron's fixed-reward-task sessions, we preferred sessions with multiple sound and contrast conditions, breaking ties by highest variance explained by the encoding model (see below), and then by trial count. Neurons with no fixed-reward session were excluded entirely from further analyses. Within the retained sessions, an area/group was only included in area- or group-level summaries if it had at least 25 good quality neurons.

#### *Serial tomography and probe tracing*

Mice were perfused transcardially and the brain was removed and post-fixed in 4% formalin-PBS (#28908, ThermoFisher) overnight and subsequently stored in PBS. Brains were then mounted into agarose and imaged using two-photon serial tomography(*98*), using a custom-built microscope with pre-processed a custom software (**https://bakingtray.mouse.vision/**)(*99*). For imaging probe tracks excitation wavelength was 780 nm, and brains were imaged in two or three channels (Red: ET570lp, Green: ET525/50m, Blue: FF01-450/70) typically at 10x10x25 µm resolution. We imaged the brains at 4x4x10 µm resolution. All images were then down sampled to 25*25*25 µm voxel size, aligned to the Allen Mouse Common Coordinate Framework (CCF)(*100*) using brainreg (in BrainGlobe), allowing us to manually track probes in atlas space using brainreg-segment (*101–103*).

To map extracellular spike activity to histological regions with high precision, we developed an automated alignment tool. Histological reference data generated with BrainGlobe were provided as per-shank tables containing structure acronyms, spatial coordinates, and variable depth indices reflecting actual trajectory lengths. Anatomical outlines and color codes for brain areas were imported from the CCF structure tree (*100*).

Spike counts from each shank were binned by depth (using probe-defined variable bin edges) and time (1 s bins) to form a depth-resolved spike histogram. Transient artifacts were suppressed by thresholding per-depth non-zero counts at the 99th percentile and interpolating missing values. The smoothed multi-unit autocorrelation (MUA) matrix was computed via pairwise cross-correlation of depth-specific spike trains. Mean firing rates per depth bin were calculated, and nonnegative matrix factorization (NNMF) with component number selected by the number of areas in the histological data. Frequency-band power profiles spanning 0.1–200 Hz were computed from depth-resolved spike train spectrograms (50 µm by 500 Hz resolution, spanning 5 min of data) and interpolated onto the primary depth axis.

Depth-specific features, including normalized firing rates, mean spike amplitudes, counts of well-isolated clusters, histological trajectory coverage, band-power intensities, and NNMF component weights, were scaled to [0,1] and concatenated into a unified feature matrix. First, the summed score of these features was used to determine the depth on the probe going from brain to liquid/air with a threshold. Depths not surviving this threshold were eliminated from the histological reference data (e.g. deep or shallow structures that were traced but not recorded from).

To align histological regions to functional domains, we first created a feature matrix F, which included normalized firing rates, mean spike amplitudes, band-power intensities, NNMF component weights, and the first principal component of the MUA matrix. Principal component analysis (PCA) on this matrix F yielded the first $n$ principal components (PC) capturing the dominant depth-dependent functional gradients, where $n$ was the component where ≥65% cumulative variance was explained. PCs were summed, and peaks in the derivative of the summed PC defined putative transition points between functional domains.

To restrain the functional domains by anatomical domains, cumulative coverage profiles for parent anatomical areas were computed by weighing each area's thickness by expected firing rates (e.g. white matter has expected firing rates of 0), with white-matter-like areas down weighted. The optimal mapping between histological and functional cumulative curves was obtained by minimizing the absolute differences in a least-squares sense. Assigned segments were refined via iterative piecewise linear interpolation of histological depths and three-dimensional track coordinates to align region boundaries with functional segments.

Aligned histology tables were converted into depth-to-area mapping tables by matching each interpolated depth to its corresponding CCF acronym and color code. Each cluster was assigned to a region by finding the minimum absolute depth difference between its depth and the depth-to-area mapping.

This tool was implemented in MATLAB R2024b within the script alignatlasdata_automated.m, available under an open-source license at github.com/EnnyvanBeest/UnitMatch/tree/main/MATLAB/Histology (Ref. (*68*)).

We then used a custom GUI to manually adjust the histology to function mapping (github.com/EnnyvanBeest/UnitMatch/blob/main/MATLAB/Histology/AlignmentCurationGUI.m, **Figure S12**).

#### *Analyses*

##### *Preprocessing*

All data were aligned to a central timeline with a time resolution of 30 Hz. For each time bin, we summed all the spikes for each neuron. We also kept a record of the average spatial position per time bin, running speed, trial number, etc.

For analysis in the spatial domain, spatial position was binned per 2.5 cm, and spike times per neuron were summed to a spike count per spatial position. Similarly, we summed the time spent at each position as 'occupancy'. Spike count and occupancy were both spatially smoothed with a Gaussian filter of width 7.5 cm, before converted to spikes/sec, and we z-scored these scores to get z(spikes/sec).

##### *Spatial modulation index*

For each neuron, full-contrast trials were split into odd (training) and even (held-out). The preferred track position was defined as the location of maximal firing in the trial-averaged, position-binned firing rate computed from the odd trials only (excluding the first 5 cm and beyond 100 cm of the track). Firing rates were then min–max normalized to [0,1] using the minimum and maximum of this same odd-trial curve, and both the odd and the held-out even-trial curves were re-centered on the preferred position and normalized with that odd-trial-derived scale, yielding a cross-validated, peak-aligned spatial response profile per neuron (±60 cm window). To get percentages, we multiplied by 100%.

Spatial Modulation Index (SMI): for each neuron, the held-out (even-trial) response at the aligned preferred position ("preferred peak") and the mean held-out response at the two sensory identical positions (±40 cm, "identical peak") were extracted. We computed a normalized Spatial Modulation Index, SMI = (preferred − identical) / (preferred + identical) to index position-specific tuning above and beyond sensory drive. Neurons (20%) were excluded if the odd- and even-trial spatial profiles were not positively correlated or when abs(SMI)>1.2, which could happen because the normalization scale is fit on the training half only (20.5% of neurons excluded), indicating the neuron's spatial tuning was not reproducible across the two independent trial halves. SMI values were then compared across brain areas using linear mixed effects (LME) models (see below) with random intercepts for mouse and session.

##### *Running correlations*

We computed cross-correlations between mean-subtracted running speed and square root transformed and mean-subtracted spike counts, with a maximum lag of 1 second. To test for significance of the (maximum) correlation, we performed a linear shift-test with a random lag 500 times to build a null-distribution of correlations. Both spike count and running speed were smoothed with a gaussian filter of 5 s width and max-min normalized for plotting purposes. We identified running bout on- and offsets and aligned spiking activity to these on- and offsets (**Figure 4d**).

##### *Bayesian decoding of position*

To determine whether position-related information was encoded in the neural population, we trained a Naïve Bayes classifier (NBc) on PCA-reduced and normalized spike count data. Spike counts were first further down sampled to 10Hz, and only trials with a minimum distance of 85cm and time points with a minimum speed of 1 cm/s were included in the analysis. Principal Component Analysis (PCA) was then applied to the time-by-neuron matrix, and the number of dimensions (k) was chosen such that the cumulative explained variance exceeded 90%.

For each session, trials were partitioned into three folds using stratified 3-fold cross-validation. Within each fold, a multinomial NBc (fitcnb.m) was trained on two-thirds of the data and tested on the remaining third using actual position bin per sample as label. To prevent an imbalance, an equal number of samples were drawn without replacement from each position bin in the training set. The number of samples per bin was set to the minimum count across bins, provided it was more than 5, otherwise the fold was omitted. The model outputs a posterior probability distribution over positions (predict.m), as well as the most likely predicted position per time point. Decoding accuracy was calculated as the proportion of positions correctly predicted as most likely.

To assess statistical significance, we computed a null distribution of accuracies by repeating the decoding procedure with position labels randomly permuted 500 times, however we kept all samples belonging to the same trial clustered, so that continuity of position within a trial was not affected when computing the null distribution. A p-value was obtained by comparing the actual decoding accuracy to this null distribution using inverse percentiles, bounded below by 1/500.

For testing differences in decoding accuracy between areas, we repeated above procedure selecting only neurons from specific areas. Although decoding accuracy increases with number of neurons, even with 10 neurons decoding accuracy was well above chance (**Figure S5a**, established by randomly subsampling *k* neurons and repeating the above procedure), hence we included areas in which at least 15 well-isolated neurons were recorded (and randomly subsampled to 15 neurons when there were more neurons). To test whether the posterior probability for the correct position was better than for the sensory identical position, we computed the log-ratio:

$$Log_{Ratio} = Log(\frac{P(correct) + \epsilon}{P(Identical) + \epsilon})$$

To see how well decoding generalizes across trial conditions, we trained a NBc for each condition (e.g. contrast = 1, audio = 1) on half of the trials and then tested the NBc on the other half of trials within condition, and trials from all other conditions.

All further statistical tests were done according to a mixed-linear model with post-hoc tests as described below.

##### *Reduced rank kernel regression*

To disentangle spatial-navigation related variables, such as sensory perception, running speed, and subjective perception of spatial position, we fitted a reduced rank kernel regression model, similar to Steinmetz et al., 2019 (*52*). This analysis only included trials on which the VR position reached at least 85cm. In this analysis, the firing rate of each neuron at timepoint $t$ is described as a linear sum of the different variables at timepoint $t$, with a time resolution of 100ms:

$$Spike\ count = Baseline + Onset + Position + Segment + Reward + Speed + Position \cdot Contrast + Position \cdot Audio + Position \cdot Contrast \cdot Audio + Position \cdot Segment + Position \cdot Segment \cdot Speed + Distance + Reward \cdot Speed,$$

where $Onset$ is the onset of the corridor at every trial (Toeplitz-lagged), $Position$ is the sensory position of the corridor (0-50cm) binned at 2.5 cm (one-hot), $Segment$ is which of the two segments of the corridor (one-hot), $Reward$ whether a reward was presented (Toeplitz-lagged), $Speed$ the running speed (Toeplitz-lagged), $Contrast$ the contrast of the visual landmarks in the trial (scalar), $Audio$ whether auditory landmarks were present or not (one-hot), and $Distance$ the physical distance traveled in the current trial (one-hot), and $Baseline$ a constant '1'. Gaussian smoothing was applied (7.5cm) was applied to the $position$ and $Distance$ terms. Toeplitz matrices were expanded over [-1.5:1.5s].

This linear model can be compactly written $Y = \beta X, \beta = (X_{train}^T X_{train} + \lambda I)^{-1} X_{train}^T Y_{train}$, with $\frac{1}{3}$ of trials held out as $test$ (in sequential blocks to avoid nonsense-correlations(*104*)). We optimized $\lambda \in [10^{-1}, 10^5]$ in nested cross-validation by minimizing $||Y_{test} - \max(\beta X_{test}, 0)||^2$. We then reduced the rank of $\beta$ via SVD, $\beta_k = U_k S_k V_k^T$, choosing $k$ to minimize the same loss. To measure performance of the model, we computed $r$: the correlation between $Y_{test}$ and $\beta_k X_{test}$.

After the example of Musall et al., 2019(*70*), we identified each variable's minimal or unique contribution by repeating above analysis with 'X' having the kernel representing the variable of interest shifted by a random amount of trials and computing the difference in $r$ with the full model. Since trials are not of equal length, we interpolate the data from the newly assigned trial into the original trial. This was repeated 500 times to obtain a p-value as the percentile of $r$ in the $r_{shift}$ distribution, with a minimum p-value of 1/500 = 0.02. We computed $r_{min}$ as the difference between $r$ and the median $r_{shift}$. Conversely, to compute the maximum contribution of a variable, we fit the model in which 'X' only included the variable of interest, and computed $r_{max}$. See below table to identify which predictors were shifted for each navigational signal of **Figure 5**.

| | *Spatial position* | *Sensory Position* | *Corridor onset* | *Visual landmarks* | *Auditory landmarks* | *Reward* | *Physical distance* | *Running speed* |
|---|---|---|---|---|---|---|---|---|
| *Onset* | | | ✓ | | | | | |
| *Position in segment* | | ✓ | | | | | | |
| *Reward* | | | | | | ✓ | | |
| *Speed* | | | | | | | | ✓ |
| *Position * Contrast* | | | | ✓ | | | | |
| *Position *Audio* | | | | | ✓ | | | |
| *Position* Contrast * Audio* | | | | ✓ | ✓ | | | |
| *Position * Segment* | ✓ | | | | | | | |
| *Position * Segment * Speed* | ✓ | | | | | | | ✓ |
| *Physical distance* | | | | | | | ✓ | |
| *Reward * Speed* | | | | | | ✓ | | ✓ |

##### *Linear mixed effects models*

Linear mixed effects (LME) models were in the form of:

$$y \sim 1 + Area + (1|Mouse) + (1|UniqueSession)$$

Random factors (1|[*variable*]) were included to account for units being recorded in the same session or mouse. We tested whether including *sex* as a variable improved the LME for every statistical test separately, which was not the case. We fit LME using MATLAB's *fitlme.* Omnibus tests of fixed effects (intercept to test whether y > 0 and area to test for differences across areas) were conducted via model comparison using *anova* on the resulting fit. Individual coefficient tests (Wald test for coefficients) were obtained with *coefTest* when fixed effects were significant and corrected for multiple comparisons using Bonferroni-Holm.

For *y* with a binomial distribution, we used the logit link type, for *y* with a gamma distribution, we used the log link type, and for *y* with a normal distribution, we used the identity link type.

##### *Dimensionality reduction (UMAP)*

We used UMAP(*105*) to find a two-dimensional non-linear embedding of the $r_{min}$ profiles. In parallel, we applied principal component analysis and projected each neuron's average loading on the UMAP.

##### *Co-selectivity*

Kernel co-selectivity was quantified using LMEs (Laplace approximation): for each predictor-target combination (excluding self-pairs), binary target activity was modelled as

$$Target \sim Predictor + Area + r + \ (1|Mouse) + (1|UniqueSession),$$

where *Predictor* is a binary indicator of anchor-kernel activity, *Area* is a categorical variable encoding the recorded brain region, *r* is the correlation between the neuron's activity and a full model (included as a continuous covariate to control for overall neural responsiveness), and mouse and session were included as random effects. Only areas with ≥25 neurons in the analysis subset and ≥10 neurons in each anchor state were included.

Adjusted odds ratios (OR) were defined as the weights given to *Predictor.* Note that similar analyses were performed where the anchor was defined as significant selectivity for a passively presented stimulus.

##### *Uniformity and precision of spatial representations*

To quantify how uniformly each brain region tiles the corridor, we first estimated each neuron's preferred position from odd-trial tuning curves (cross-validated against even trials, full-contrast trials only). Neurons with a tied maximum were excluded. To avoid a non-spatially specific transient at the start of the corridor being misread as a 'preferred position near onset', the first 5 cm of the corridor were excluded before computing preferred positions (except for the histograms in **Figure 6a-b**).

For each group of neurons, we repeatedly (50 times) drew a random subsample of N (200 for brain groups, 100 for individual regions) neurons without replacement from the pooled population, and computed the Shannon entropy of the resulting distribution of preferred positions across corridor bins:

$$H_{norm} = \ -\sum_i p_i \frac{\log_2(p_i)}{\log_2(\,Npos)},$$

Where $P_i$ is the proportion of neurons whose preferred position falls in bin *i* and *Npos* is the total number of position bins. A value of 0 indicates all neurons share the same preferred location; a value of 1 indicates perfectly uniform coverage of the corridor.

Entropy was then normalized by the maximum possible entropy (drawing N neurons uniformly over the corridor) (*74*), and expressed as a percentage, so that 100% represents the most uniform distribution as can be expected. Values below 100% indicate more clustering relative to maximum possible entropy. Because repeated subsample draws from the same underlying neurons are not independent observations, we tested whether uniformity differed across groups of regions with a permutation test rather than treating each draw as an independent sample: we computed a Kruskal-Wallis $\chi^2$ statistic across groups on the 50-draws-per-group uniformity values, then built a null distribution (1000 permutations) by randomly reassigning neurons to groups (preserving each group's pool size) and repeating the identical draw-and-metric procedure on the shuffled assignment. The reported p-value is the fraction of null $\chi^2$ values at least as large as the observed one.

Using the same matched-size random subsamples as above, we computed the Pearson correlation between odd- and even-trial spatial tuning curves at every pair of position bins, giving a 39 × 39 population-vector correlation matrix per subsample (39 = the number of 2.5 cm bins spanning the trimmed 5-100 cm corridor) (*75*, *76*). Averaging this matrix's off-diagonals at each spatial offset gives a correlation-vs-offset decay curve. We fit a straight line to this decay curve over offsets from 0 to 14 cm (the extent of one "sensory segment" before the next landmark) after first rescaling the decay curve to [0,1] between that subsample's own floor (mean correlation beyond 30 cm) and peak (correlation at offset 0) correlation, to express precision as a fraction of each area's own achievable correlation range rather than raw r per cm. Higher values ($cm^{-1}$) indicate a more precise spatial code. Values were averaged across the 50 draws, and the same permutation procedure described above was used to test for a difference in precision across groups of regions.

Finally, for each neuron with a defined preferred position, we computed its signed offset from its session's fixed reward location and, separately, its signed offset from whichever of the four visual landmarks (at 20, 40, 60 and 80 cm) was numerically nearest to it. Within each group of regions, these per-neuron offsets were binned using edges shared across every group and reference point (5 cm bins spanning ±102.5 cm around zero), from which we derived the fraction of that group's neurons falling in the bin with the most preferred responses ("peak fraction"). We tested whether each group's peak fraction was more extreme than expected by chance using a two-sided permutation test: a null distribution was built by repeatedly (2000 times) drawing a same-size random subset, without replacement, from the pooled offsets of every group being compared for that reference point (including the group itself) and recomputing the peak fraction over the same bin edges.

As a robustness check, we repeated the above uniformity and precision analyses after regressing out each neuron's trial-by-trial dependence on running speed from its firing rate before recomputing tuning curves, to confirm that the reported differences in uniformity and precision across groups of regions were not confounded by systematic differences in running speed at different corridor positions.

### Figures S1 to S12

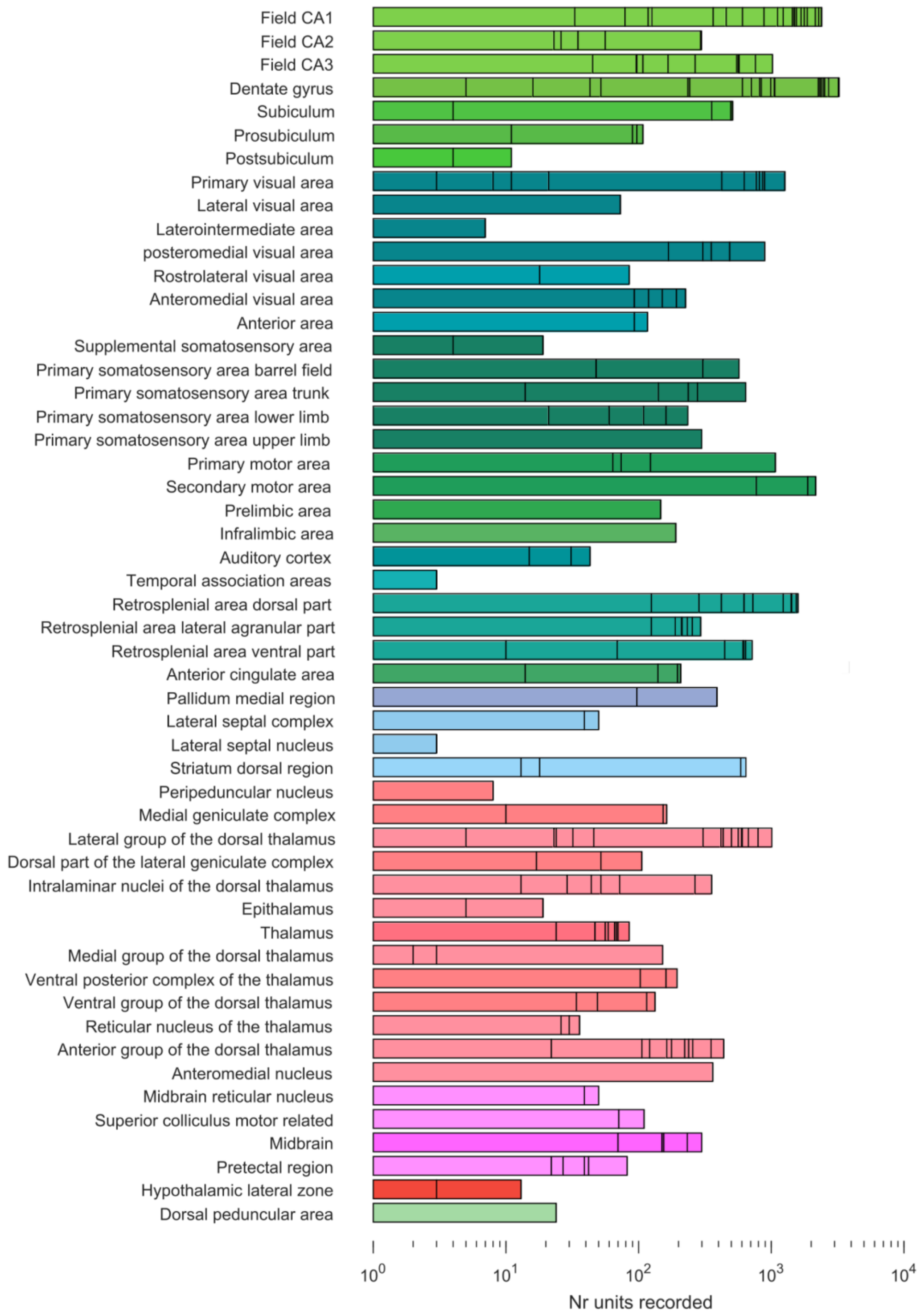


***Figure S1.*** *Number of well-isolated single neurons per brain region.*

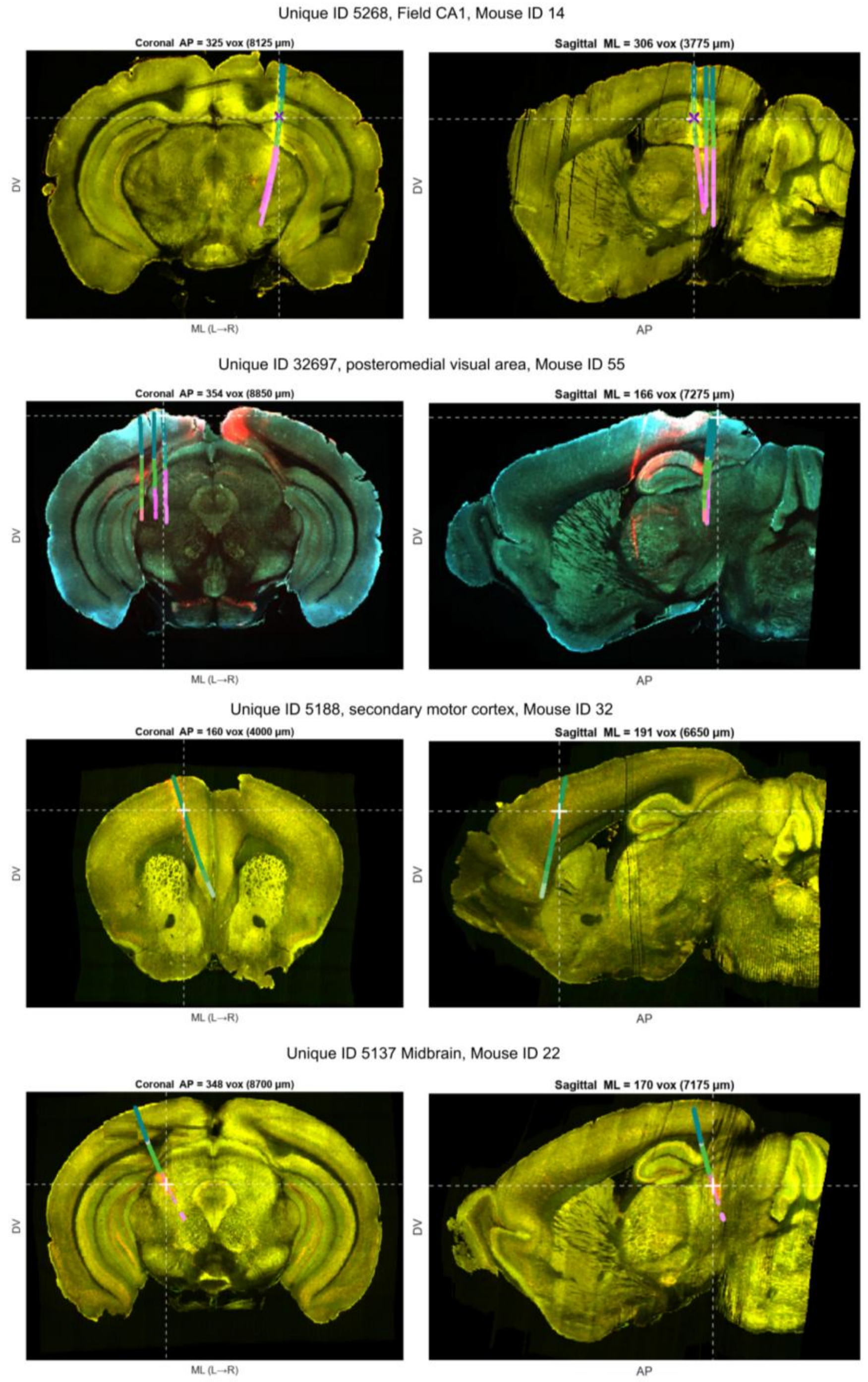


***Figure S2.*** *Histology for example neurons in **Figure 1**. Neuropixels shanks depicted on top of the brain, with channels colored by the brain region. Top to bottom: i to iv.*

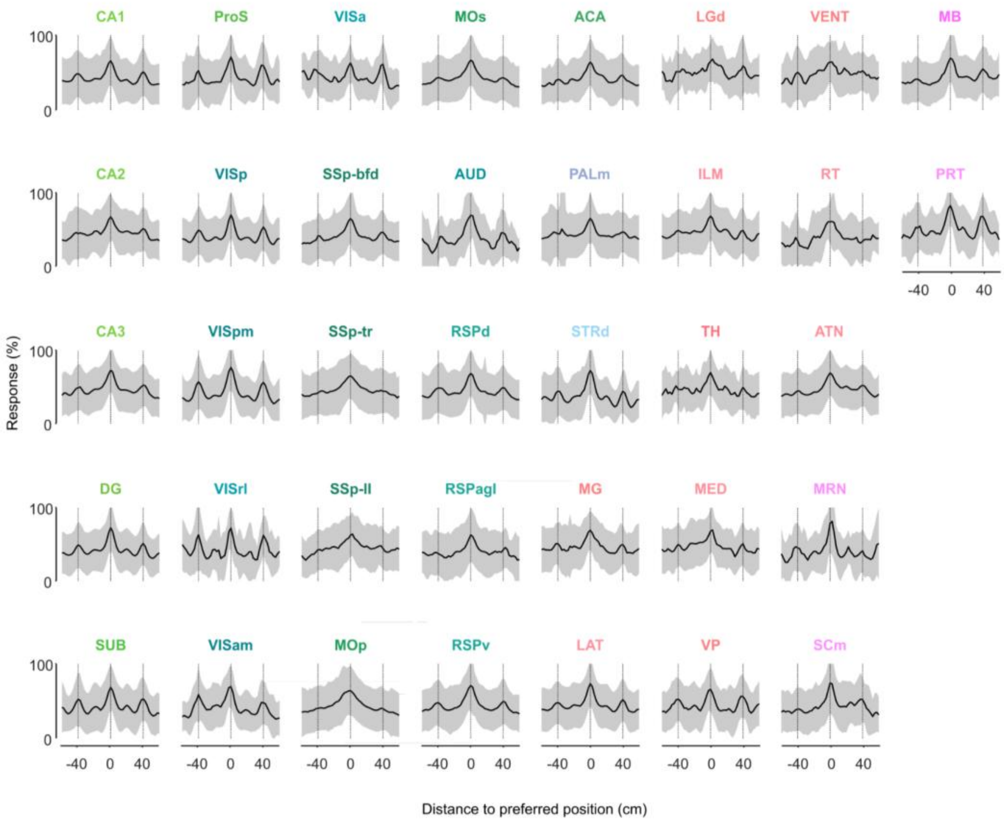


***Figure S3*. *Normalized activity (mean ± s.d. % across neurons)*** *as a function of distance to preferred position (measured in held-out trials). Same as* ***Figure 1****q-x for all regions separated.*

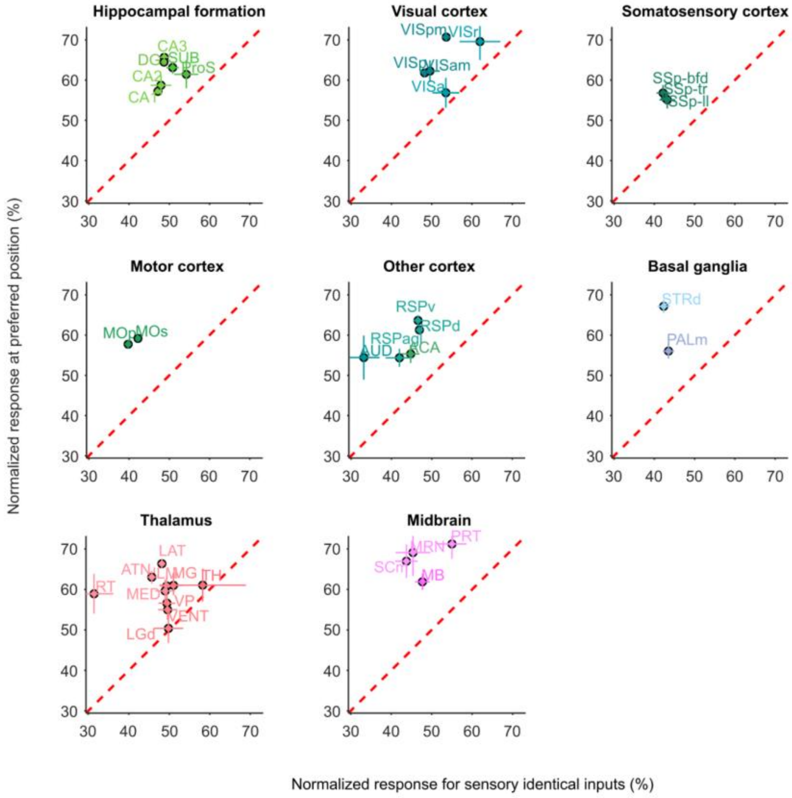


***Figure S4. Response at preferred versus sensory identical position.***

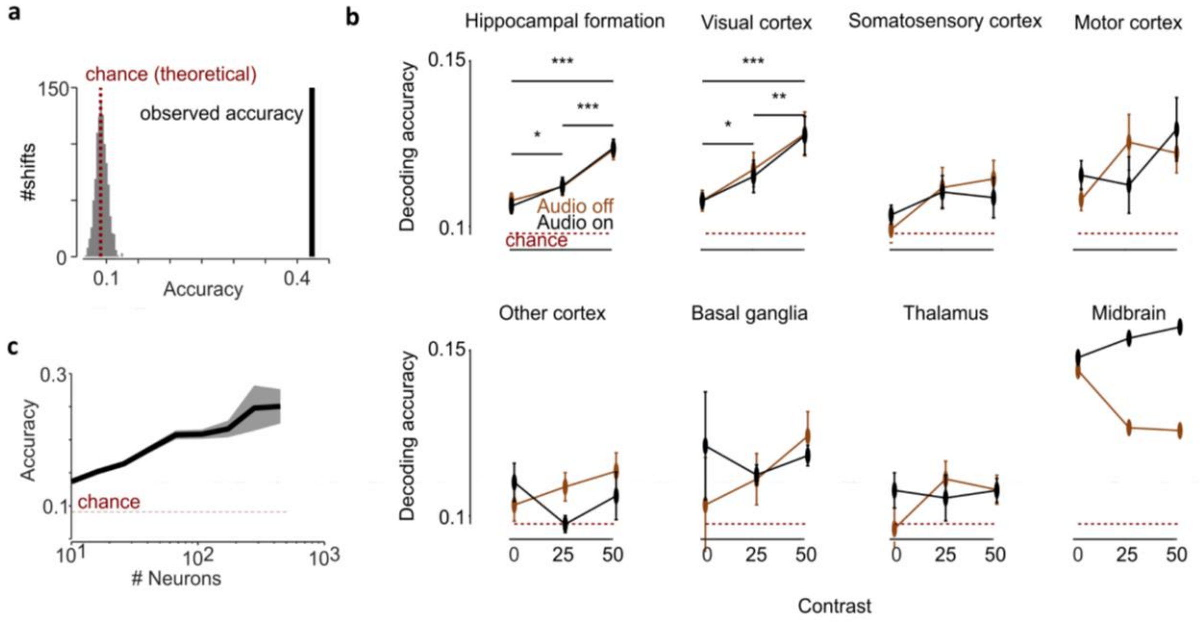


***Figure S5. Bayesian decoding. a)*** *Shows an example session of observed accuracy (black line) of position decoding against a theoretical chance level (red dotted line) and a null distribution of accuracies (in gray) observed when we linearly shifted position in time.* ***b)*** *Decoding accuracy (mean ± s.e. across sessions) of position increases with the number of neurons included, but is already above chance with as little as ten neurons.* ***c)*** *Decoding accuracy as a function of visual contrast (x-axis) for audio off (brown) and audio on (black) trials, split up per brain region. Holm-corrected significance after established main effect.*, p < 0.05; **, p<0.01; ***, p<0.001.*

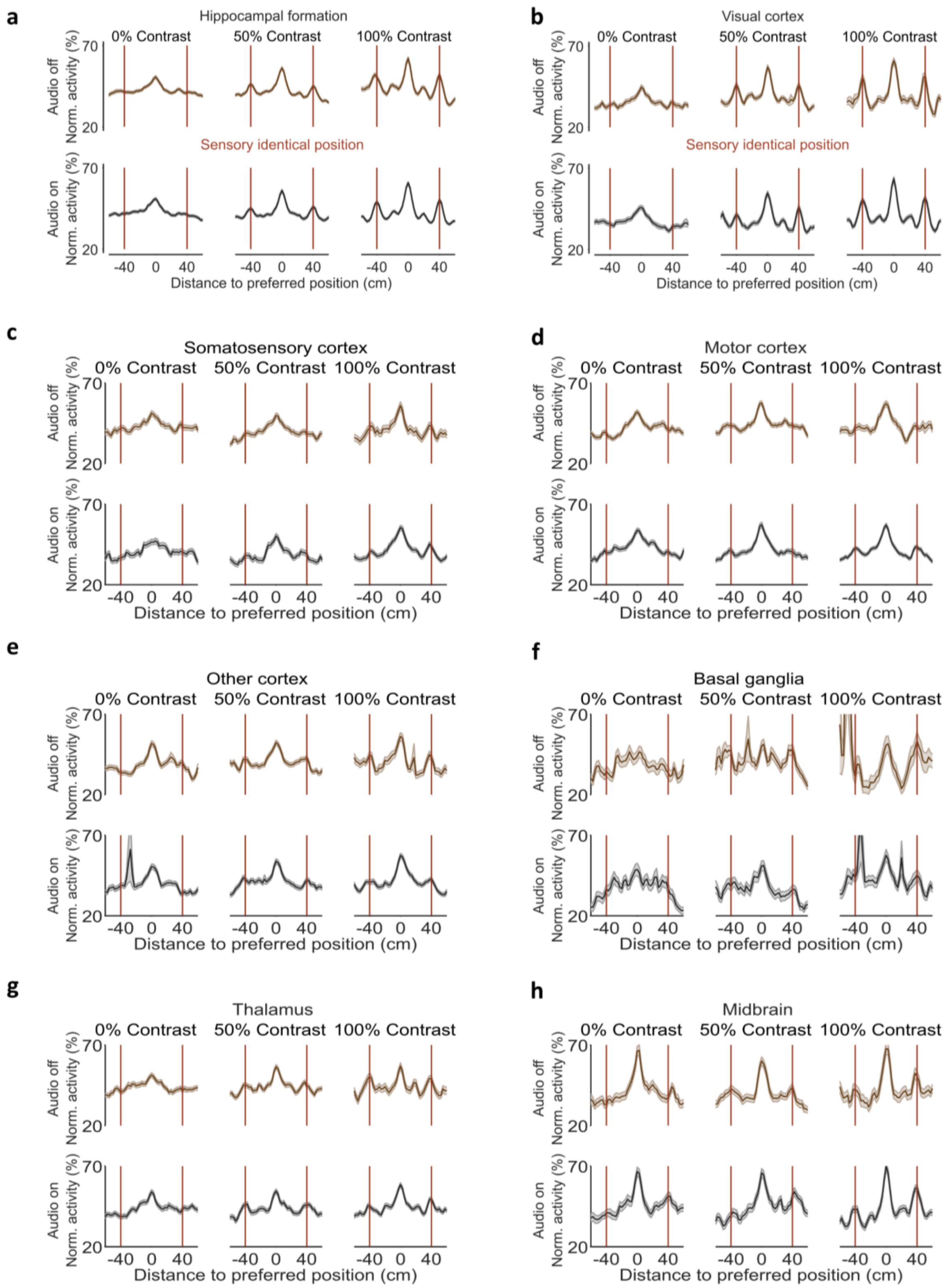


***Figure S6. Normalized activity*** *(mean ± s.e. across neurons) as a function of distance to preferred position (0-tick), separated for different trial-conditions. Red line indicates the identical position(s). Preferred position was determined in held-out full contrast audio on trials. For different brain groups.* ***a)*** *hippocampal formation,* ***b)*** *visual cortex,* ***c)*** *Somatosensory cortex,* ***d)*** *motor cortex,* ***e)*** *other cortex,* ***f)*** *basal ganglia,* ***g)*** *Thalamus,* ***h)*** *Midbrain.*

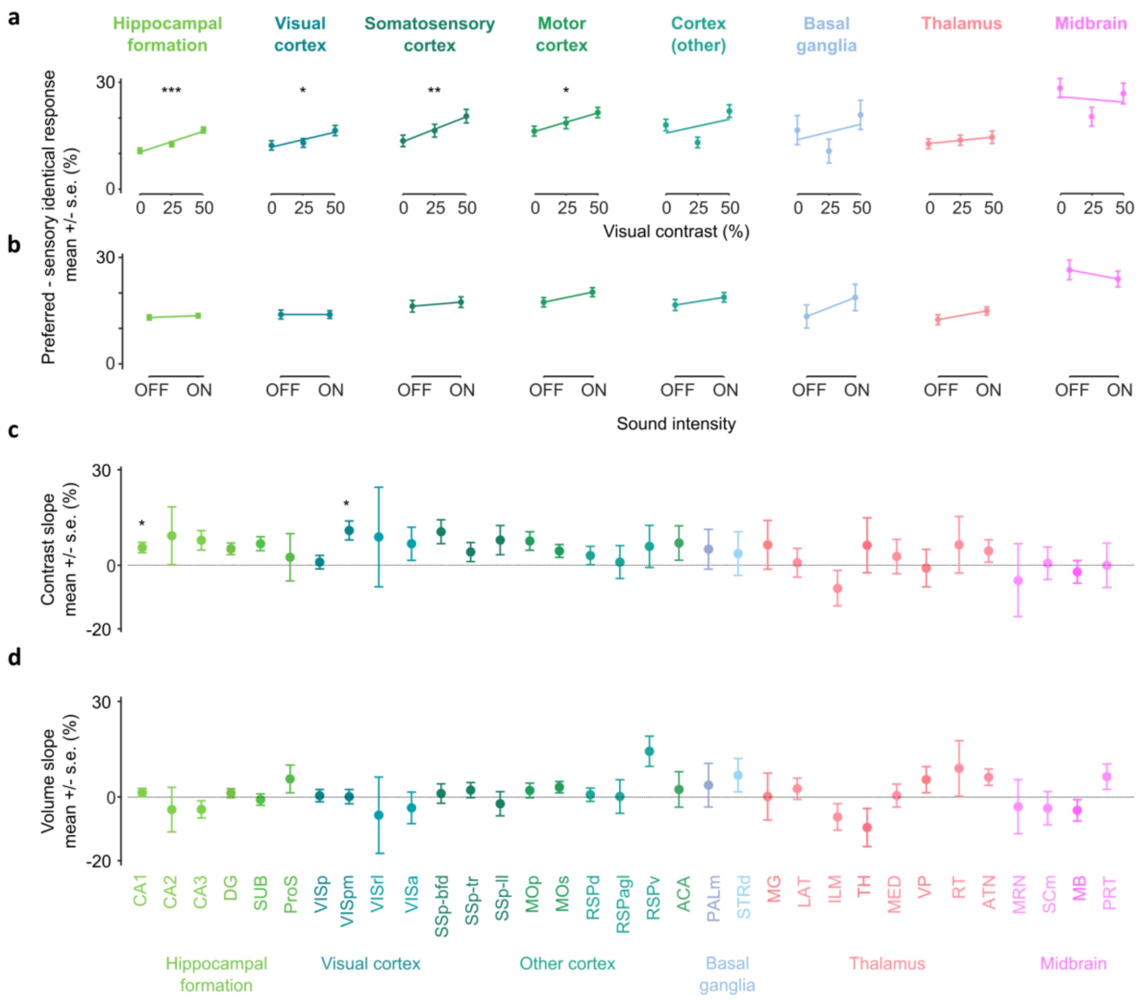


***Figure S7. Anchoring to visual and auditory landmarks. a)*** *Difference (mean ± s.e.) between response at preferred position versus sensory identical positions for visual contrast conditions, averaged across sound intensity conditions. Line is a linear fit to the data. ***, p<0.001; **, p<0.01; *, p<0.05 for significant effect of contrast (LME).* ***b)*** *Same as a) but for different sound intensity conditions, averaged across visual contrast conditions.* ***c)*** *Slope (mean ± s.e. across neurons) of the contrast fit as illustrated in a).* ***d)*** *same as c) for volume. No significant effect of brain region (LME).*

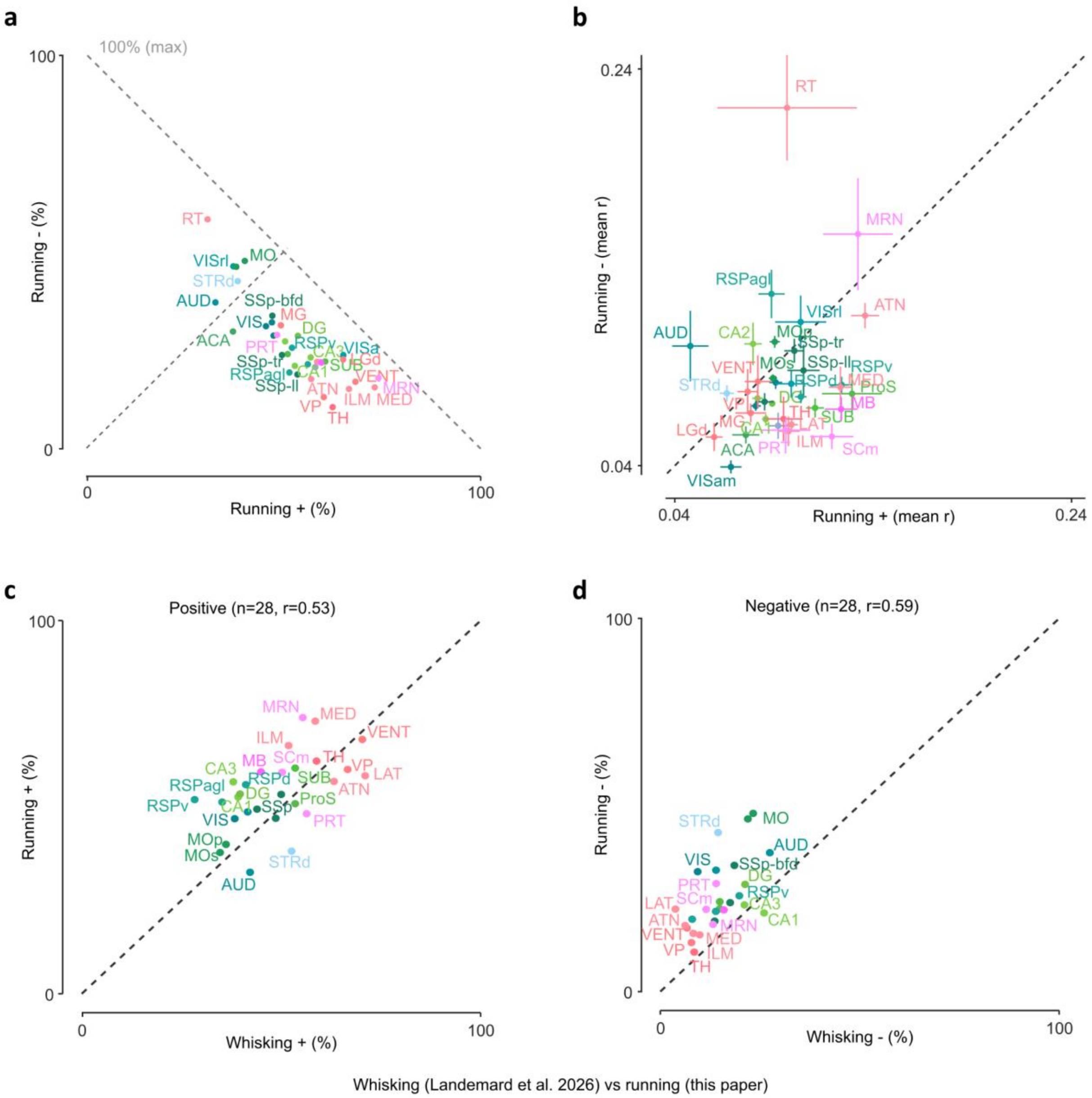


***Figure S8. Brainwide correlations with running**. **a)** Summary of **Figure 4e;** percentage of positively versus negatively correlating neurons per brain region. **b)** mean correlation for positive and negatively correlating neurons per brain region. **c-d)** Comparison with whisking(71): percentage of positively (c) and negatively (d) correlating neurons.*

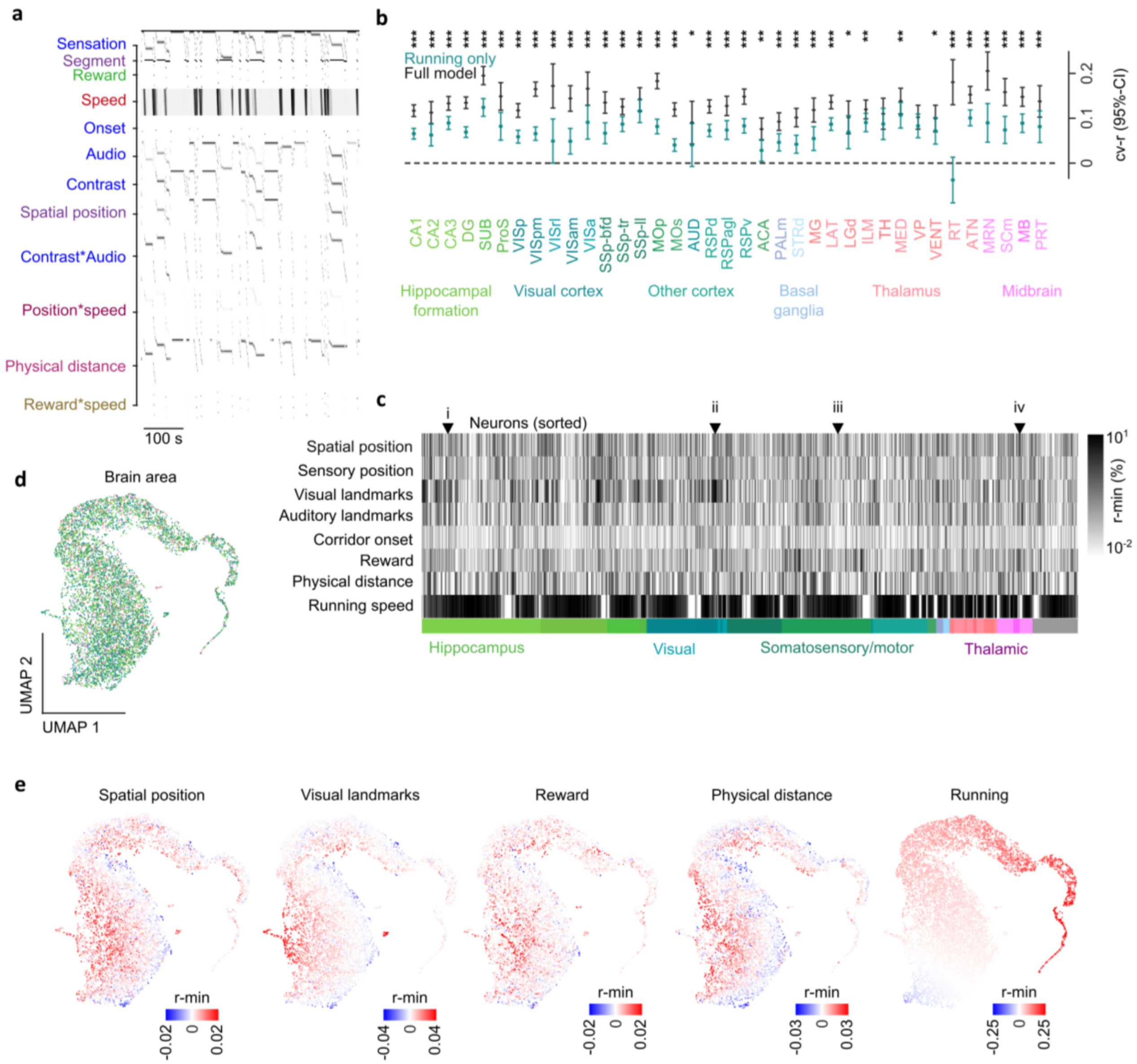


***Figure S9. General linear model of a virtual reality corridor. Related to Figure 5. a)*** *Design matrix for example session, showing the different variables over time, together explaining events in the virtual corridor.* ***b)*** *r-min profiles across predictors per neuron. Neurons were sorted first by area and then by hierarchical clustering. Example neurons are indicated.* ***c)*** *Cross-validated correlation (r) estimates (95% confidence interval) for the full model (black) and a model including only running as a predictor (aqua). Estimated from an LME, ***, p < 0.001; **, p < 0.01; *, p < 0.05 for post-hoc comparisons full versus running only model.* ***d)*** *UMAP projection (minimal distance: 0.5, number of neighbors: 5; varying these parameters gave overall the same results) of r-min profiles, colored by different predictors: brain area (same colormap as underneath b);* ***e)*** *UMAP colored by spatial position, visual landmarks, rewards, physical distance, and running. All show r-min contributions in red-blue color scale. Note that running r-min values are in an opposite axis than other navigational variables.*

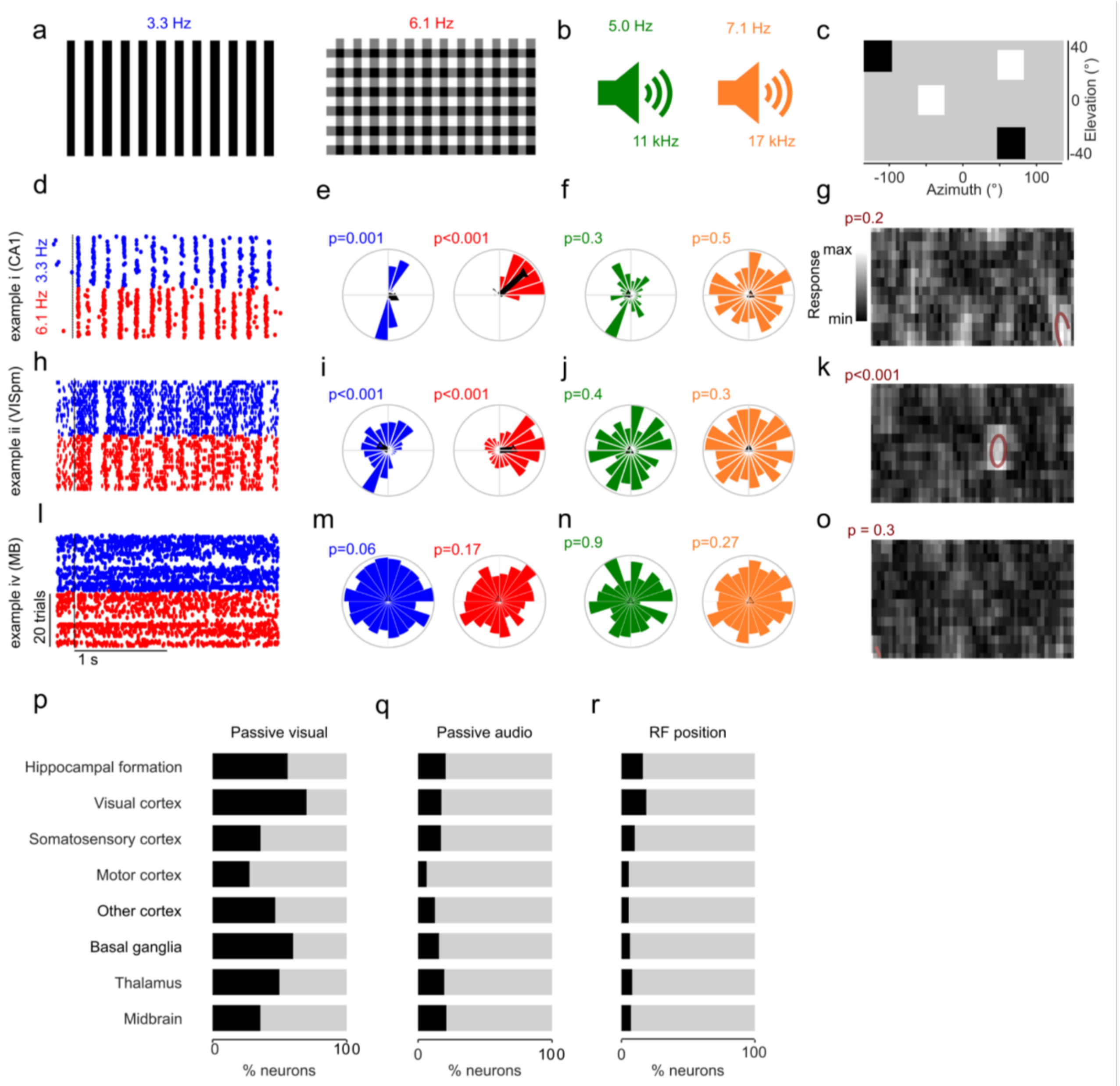


***Figure S10. Passive sensory tuning predicts navigational signals. a)*** *Full screen vertical grating (left) and plaid (right) were presented one by one at random in some recordings after mice navigated in the virtual corridor. Stimuli were presented at specific frequencies to allow for frequency tagging of neural responses.* ***b)*** *A cosine-modulated pure tone of either 11 or 17 kHz (randomized) was presented simultaneously with the presentation of the full screen plaid or grating. Tones were also played at a specific frequency to allow for frequency tagging.* ***c)*** *Receptive fields were mapped with sparse noise stimuli.* ***d)*** *Spikes as a function of time after stimulus onset for an example neuron in CA1 (same a* ***Figure 1e****). Trials are sorted by condition of the visual grating.* ***e)*** *Polar plots showing the (normalized) number of spikes per phase of the frequency the respective visual stimulus was shown. Black bar shows the circular mean and mean vector length. p-values: Rayleigh test for non-uniformity of circular data.* ***f)*** *Same for the auditory stimuli.* ***g)*** *Normalized responses in azimuth and elevation to sparse noise stimuli. Red ellipses are the best fit of a receptive field (see methods), and a p-value of the response relative to a shuffled null-distribution.* ***h-k)*** *Same for example neuron in VISpm (same a* ***Figure 1f****).* ***l-o)*** *Same for example neuron in MB (same a* ***Figure 1b****).* ***p)*** *Percentage of neurons per brain group significantly tuned to passive visual stimuli.* ***q)*** *Percentage of neurons per brain group significantly tuned to passive auditory stimuli.* ***r)*** *Percentage of neurons per brain group with significant receptive fields.*

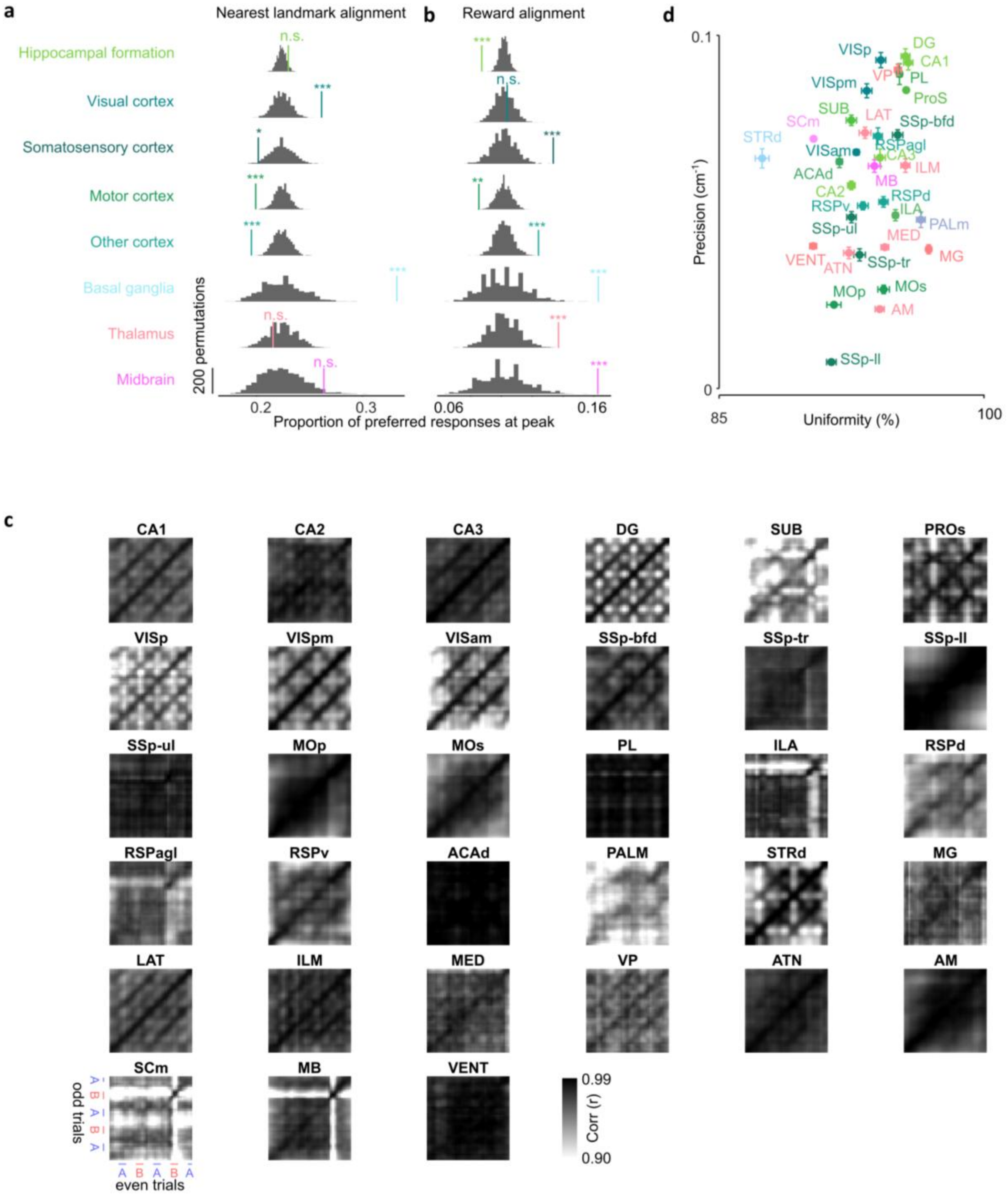


***Figure S11. Population representations: clustering and precision (related to Figure 6).*** *a) Permutation test for clustering of preferred positions around the nearest of the four corridor landmarks, per group of regions. Gray histograms show the null distribution obtained by permuting neuron-to-group assignment, colored ticks mark each group's observed proportion of preferred responses falling in its single most populated offset bin, with two-sided permutation significance (n.s., not significant; *, **, *** = p < 0.05, 0.01, 0.001). b) As in (a), for alignment to reward instead of the nearest landmark. c) Population-vector correlation matrices (odd- vs. even-trial responses) for each individual Allen CCF area with sufficient neurons (N = 100), corresponding to Figure 6c computed at the level of individual regions rather than the eight groups of regions. d) Precision (cm$^{-1}$, as in Figure 6f) against uniformity (%) for the same individual areas as in (c). Error bars, mean ± s.e.m. across draws.*

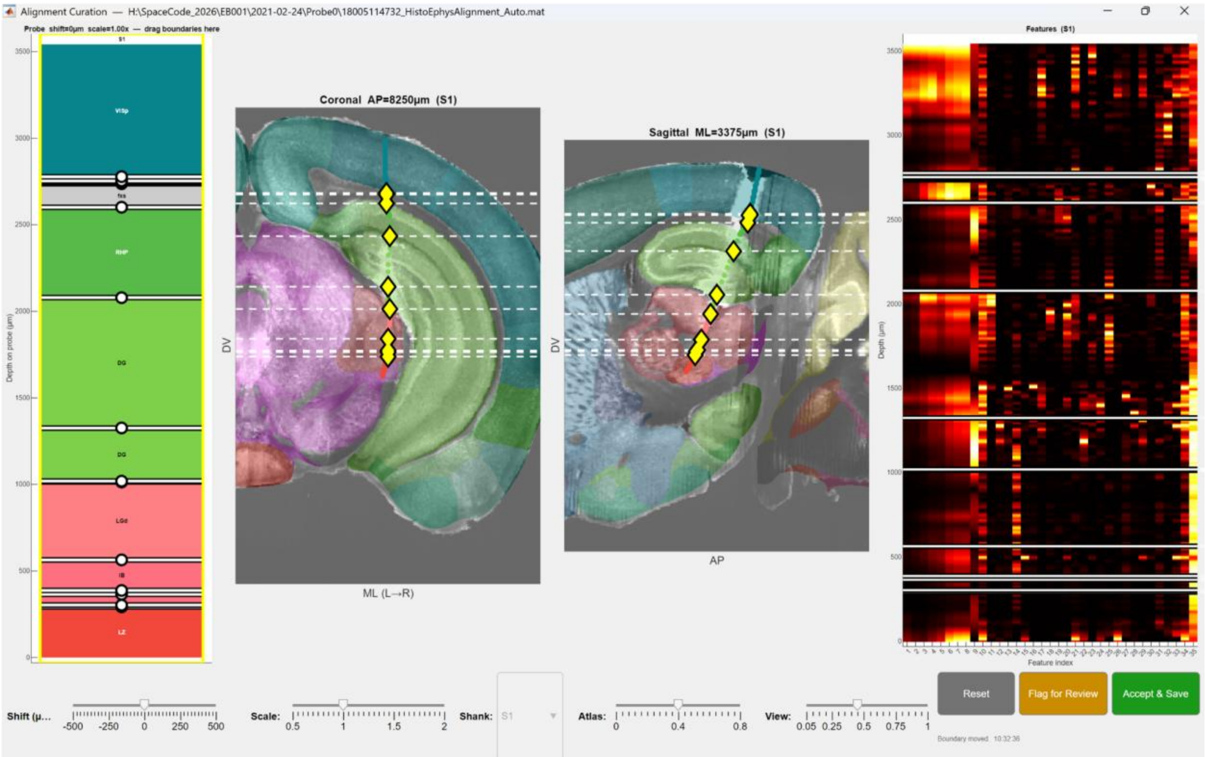


***Figure S12*. *Histology alignment GUI for manual curation.*** *Left: areas at different depths on the probes. Borders can be dragged to better align to the histological and functional data of individual probe insertions. Middle: coronal and sagittal view of probe trajectory, with colors corresponding to the brain area as indicated on the probe view (left), and yellow diamonds indicating the borders. Right: Functional electrophysiological markers (columns) as a function of depth on the probe. See the methods section for details on the specific markers. White lines indicate area borders, which are expected where electrophysiological marker profiles change in depth.*

### Tables S1 to S2

| ID | Sex | Probe | Implant | Bonus reward | Age (surgery) |
|---|---|---|---|---|---|
| 01 | M | Phase 3B | Acute | No | 70 |
| 03 | M | Phase 3B | Acute | No | 78 |
| 06 | M | Phase 3B | Acute | No | 133 |
| 07 | M | Phase 3B | Acute | No | 68 |
| 09 | F | Phase 3B | Acute | No | 79 |
| 10 | F | Phase 3B | Acute | No | 82 |
| 11 | F | Phase 3B | Acute | No | 82 |
| 14 | M | 2x 4 shanks | Chronic | No | 84 |
| 15 | M | 2x 4 shanks | Chronic | No | 115 |
| 17 | F | Phase 3B | Acute | No | 66 |
| 19 | M | 2X 4 shanks | Chronic | No | 72 |
| 22 | F | Phase 3B | Acute | Yes | 176 |
| 27 | F | Phase 3B | Acute | Yes | 69 |
| 29 | M | Phase 3B | Acute | Yes | 133 |
| 32 | F | Phase 3B | Acute | Yes | 87 |
| 33 | F | Phase 3B | Acute | Yes | 212 |
| 34 | F | Phase 3B | Acute | Yes | 213 |
| 35 | F | Phase 3B | Acute | Yes | 218 |
| 36 | M | 2x 4 shanks | Chronic | No | 75 |
| 37 | M | 2x 4 shanks | Chronic | No | 75 |
| 38 | M | Phase 3B | Acute | Yes | 50 |
| 39 | M | Phase 3B | Acute | Yes | 56 |
| 48 | F | 2x 4 shanks | Chronic | Yes | 95 |
| 49 | F | 2x 4 shanks | Chronic | Yes | 98 |
| 51 | M | 2x 4 shanks | Chronic | Yes | 158 |
| 53 | M | 2x 4 shanks | Chronic | Yes | 51 |
| 54 | M | 2x 4 shanks | Chronic | Yes | 79 |
| 55 | M | 2x 4 shanks | Chronic | Yes | 84 |

***Table S1.*** *Mouse inclusion table. Bonus reward means mice were given a double reward if they slowed down within 6cm of the reward position.*

| Area | 1 | 3 | 6 | 7 | 9 | 10 | 11 | 14 | 15 | 17 | 19 | 22 | 27 | 29 | 32 | 33 | 34 | 35 | 36 | 37 | 38 | 39 | 48 | 49 | 51 | 53 | 54 | 55 | Total |
|---|---|---|---|---|---|---|---|---|---|---|---|---|---|---|---|---|---|---|---|---|---|---|---|---|---|---|---|---|---|
| TH | 24 | | | | | | | | | 23 | | | | | 9 | | 3 | 7 | | | | 2 | | 2 | | 15 | | | 85 |
| MG | 10 | | | | | | | | | | | | | | | | | | | | | | | | | 143 | 10 | | 163 |
| LAT | 5 | | 18 | | | | | 1 | | | | | 8 | 14 | | 260 | 111 | 18 | | | 67 | 62 | 33 | 8 | 67 | 124 | 214 | | 1010 |
| DG | 5 | 11 | 27 | | 9 | | | 183 | | 8 | 364 | | 101 | 109 | 23 | 150 | 64 | 10 | 1200 | | 60 | 65 | 73 | 63 | 177 | 499 | 41 | | 3242 |
| CA1 | 1 | 32 | 46 | | 39 | | 8 | 239 | | 93 | 150 | | 273 | 233 | 115 | 216 | 38 | 15 | 51 | 129 | 96 | 94 | 283 | 126 | 115 | | | 8 | 2400 |
| SUB | 4 | | | | | | | 351 | | | | 142 | | | | | | | | | | | | | | | 15 | | 512 |
| VISp | 3 | | | | | 5 | 3 | | | 10 | 403 | 202 | 146 | 41 | 1 | 48 | | 29 | | 377 | | | | | | | | | 1268 |
| MED | | 2 | 1 | | | | | | | | | | | | | | | | | | | | | | | 149 | | | 152 |
| ILM | | 13 | 16 | | | | | | | | | | | | 15 | | 8 | | | | | 20 | | | | 194 | 89 | | 355 |
| RSPd | | 125 | 160 | | | | | | | 136 | | 203 | 1 | 102 | | 503 | 181 | | | | 2 | 12 | 118 | 51 | | | | | 1594 |
| STRd | | | | 13 | | 5 | | | 56 | | | | | | | | | | | 570 | | | | | | | | | 644 |
| SSp-ll | | | | 21 | | 39 | | | 74 | | | | | | | | | 49 | | | | | | | | | | 52 | 235 |
| VISam | | | | | 93 | | | | | | | | | | | 26 | 32 | | | | | 42 | | | 33 | | | | 226 |
| SSp-tr | | | | | | 14 | | | | 127 | | | 96 | | | | | 41 | | | | | | | | | | 364 | 642 |
| AUD | | | | | | 15 | | | | 16 | | | | | 12 | | | | | | | | | | | | | | 43 |
| SSp-bfd | | | | | | 48 | | | 266 | | | | | | | | | | | 257 | | | | | | | | | 571 |
| VISrl | | | | | | 1 | 17 | | | | | | | | | | | | | 67 | | | | | | | | | 85 |
| RT | | | | | | | | | | 26 | | | | 4 | 6 | | | | | | | | | | | | | | 36 |
| ATN | | | | | | | | | | 22 | | | 84 | 15 | | 42 | 14 | 46 | | | | 16 | | | | 17 | 96 | 86 | 438 |
| CA3 | | | | | | | | | | 45 | | | 51 | 1 | 11 | | | 59 | 100 | 280 | | 1 | 10 | 9 | | 5 | 188 | 261 | 1021 |
| CA2 | | | | | | | | | | 23 | | | 3 | 9 | | | | | 21 | 237 | | | | | | | | 5 | 298 |
| RSPagl | | | | | | | | | | 125 | | | 64 | 21 | 3 | | | | 20 | | | | 21 | 40 | | | | | 294 |
| VP | | | | | | | | | | 103 | | | | | | | | | | | | | | 58 | | 34 | | | 195 |
| RSPv | | | | | | | | | | 10 | | 59 | | | | 377 | 167 | | | | 5 | 23 | | | | | | 78 | 719 |
| MRN | | | | | | | | | | | | 39 | | | | | | | | | | | | | | 11 | | | 50 |
| SCm | | | | | | | | | | | | 71 | | | | | | | | | | | | | | | | 39 | 110 |
| MB | | | | | | | | | | | | 70 | | | 80 | | 5 | | | | | | | | | 78 | | 66 | 299 |
| ProS | | | | | | | | | | | | 11 | | | 79 | | | 7 | | | | | | | | 11 | | | 108 |
| PALm | | | | | | | | | | | | 97 | | | | 293 | | | | | | | | | | | | | 390 |
| ACA | | | | | | | | | | | | 14 | | | 126 | 57 | | | | | 11 | | | | | | | | 208 |
| MOp | | | | | | | | | | | | | | 64 | 10 | | | | 49 | | | | | | 952 | | | | 1075 |
| MOs | | | | | | | | | | | | | | | 772 | | | | | 1114 | | | | | 279 | | | | 2165 |
| PRT | | | | | | | | | | | | | | | 22 | | | 5 | | | | | | | 12 | 3 | | 40 | 82 |
| VISpm | | | | | | | | | | | | | | | 168 | 137 | 48 | | | | 132 | 1 | | | | | | 407 | 893 |
| LSX | | | | | | | | | | | | | | | | 39 | | | | | 11 | | | | | | | | 50 |
| VENT | | | | | | | | | | | | | | | | | | | 34 | | | 15 | | | | 66 | 18 | | 133 |
| LGd | | | | | | | | | | | | | | | | | | | 17 | | | | 35 | 54 | | | | | 106 |
| VISa | | | | | | | | | | | | | | | | | | | | | | | 93 | | 24 | | | | 117 |
| Total | 52 | 183 | 268 | 34 | 141 | 127 | 28 | 774 | 396 | 767 | 917 | 908 | 827 | 613 | 1452 | 2148 | 671 | 286 | 1492 | 3031 | 384 | 353 | 666 | 411 | 1659 | 1349 | 671 | 1406 | 22014 |

***Table S2.*** *Number of well-isolated neurons per mouse and per area. Abbreviations are consistent with the Allen Mouse Common coordinate atlas.*
